# Self-extending DNA recording with stochastic and programmed writing

**DOI:** 10.64898/2026.10.14.759016

**Authors:** Zhihui Zhang, Yingyi Li, Yifang Peng, Rongsong Ling, Kang Li, Maosheng Cheng, Yixin Cheng, Caihua Zhang, Xiaochen Wang, Shuang Chen, Zhenyu He, Demeng Chen

## Abstract

Recording cellular histories requires continued information accumulation while retaining earlier records. Infinite Printer couples DNA self-extension to regeneration of a writable terminal context. A single self-targeting prime-editing guide directs repeat extension, while base editing diversifies the accessible sequence in situ. A separate configuration uses auxiliary guides for specified payload writing. After three rounds of payload input, 4,436 distinct complete records contained identities from all three pools. Extended AC/AT records retained shared upstream marks, and complete-sequence readout distinguished endpoints sharing initial prefixes. Joint single-cell readout recovered 1,549 records and 7,669 combinations across 12,302 transcriptome-linked cell labels; spatial readout linked records to 11,330 capture spots. These results establish a resource for self-extending stochastic and programmed DNA recording with joint cellular and tissue readout, providing a foundation for future lineage applications.

## Introduction

Cellular histories shape development and tumour evolution, yet an endpoint molecular profile provides only an indirect account of how a cell reached its present state [1, 2, 3]. Genetic recorders store heritable changes in DNA, allowing cellular history to be read alongside cellular phenotypes. Genome-editing recorders have traced developmental relationships [4, 5, 6, 7, 8], and joint lineage and gene-expression readouts have been applied to zebrafish and mammalian development [6, 9, 10, 11] and experimental tumour dissemination [12], complemented by in situ and sequencing-based spatial readouts [13, 14]. Recording across large cell populations over prolonged mammalian development, however, requires more than a diverse set of endpoint states: the substrate must continue to acquire new heritable information while retaining access to information written earlier, with writing kinetics suited to the timescale of lineage branching [6, 15]. We therefore investigate self-extending barcodes as a design strategy for expanding the recording substrate to accommodate further information while preserving the readability of accumulated sequence.

Existing recording architectures use different substrate designs to support continued writing and organize accumulating sequence information. Target arrays and homing guide RNAs accumulate heritable changes, but recording can exhaust addressable sites or disrupt continued targeting [4, 5, 6, 7, 9, 16, 17]. For ordered recording, CHYRON combines a homing guide with terminal deoxynucleotidyl transferase to support insertional recording at a single site [18]. DNA Typewriter writes ordered symbols into a designed tandem substrate [15], whereas peCHYRON uses propagating prime-editing insertions to extend an ordered record [19]. PEtracer installs predefined lineage marks at multiple integrated cassettes and couples recording to sequencing and imaging [20]. DOMINO, by contrast, implements sequential base-editing cascades within predesigned repeat arrays without extending the DNA substrate, including designs in which a single guide activates successive binding sites [21]. Building on these strategies, we asked whether substrate growth and character generation could be assigned to separate components within one recording architecture: a single self-targeting guide design would sustain extension, while a separate editor would diversify the accessible sequence in situ without specifying a different insertion template for every character state.

Our self-targeting prime-editing guide RNA (stpegRNA) contains repeat-complementary arms designed to retain a compatible terminal target after extension [22]. AncBE4max diversifies accessible sequence [23]. In the normalized record orientation, earlier sequence remains towards the 5*^′^* end while new sequence is added near the regenerated 3*^′^* terminus, providing a design basis for relative preservation of earlier content. Auxiliary pegRNAs provide an alternative mode for specified payload writing.

Here we characterize repeat extension, in situ diversification and programmed writing, retaining complete sequences and assay-specific support. Matched representations and models examine the information and candidate relationships supported by those records. Single-cell and spatial transcriptomic workflows link recording to gene-expression profiles and tissue coordinates.

## Results

### A renewable substrate for stochastic writing

Infinite Printer–Stochastic Writing (IP-SW) combines an extendable AC-repeat substrate with local C-to-T diversification (Figure 1a). A single stpegRNA-AC design retains a compatible substrate after extension (Figure 1b,c; Figure S1). In the design model, RTT-specified AC addition preserves the GCA–GGG terminal context while earlier sequence remains upstream (Figure 1c,d). AncBE4max converts accessible AC units to AT, separating character diversification from substrate renewal. Transcript-linked stable lines used doxycycline-inducible PE, whereas bulk DNA experiments used transient expression (Figure 1e,f; STAR Methods).

**Figure 1.**
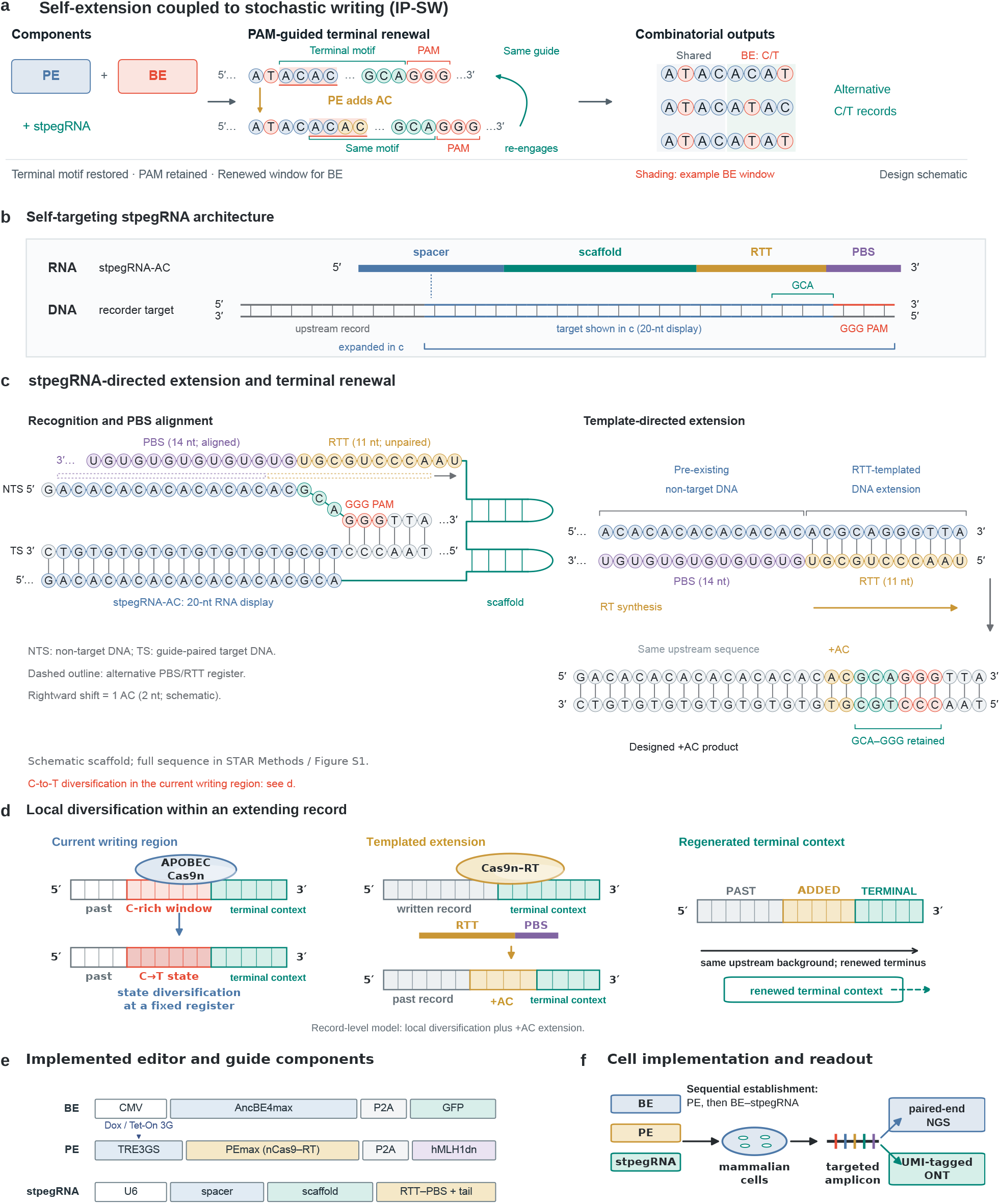
Self-extension coupled to stochastic writing. a, IP-SW separates PE-directed substrate extension from BE-directed C/T diversification. b, Guide domains and antiparallel DNA target; the displayed 20-nt target includes GCA and precedes the GGG PAM. c, Design model of extension and terminal renewal. The 14-nt PBS and 11-nt RTT occupy one RNA strand. The dashed alternative register indicates a one-AC shift, not a measured transition. RTT specifies ACGCAGGGTTA in DNA orientation, including an additional AC unit. The designed product retains upstream sequence and GCA–GGG. RNA uses U; DNA uses T. d, Record-level relationships between diversification, extension and terminal renewal; the added unit is ochre and terminal context teal. Relative retention refers to the position of earlier sequence upstream of the renewed terminus. e, Inducible PEmax-P2AhMLH1dn and AncBE4max-P2A-GFP expression cassettes used for transcript-linked stable implementation. U6 drives stpegRNA. f, Sequential establishment of PE and BE–stpegRNA stable lines and targeted readout. Bulk DNA experiments used transient editors; IP-PW omitted BE. The schematics depict the proposed design mechanism. Complete sequences are in Figure S1 and construct resources; experimental conditions are in STAR Methods.

### Programmed payload accumulation in an extending register

We next asked whether the renewable recorder substrate could support repeated writing of specified sequences. We therefore developed a base-editor-free programmed-writing configuration, Infinite Printer–Programmed Writing (IP-PW), combining a CCA-extending stpegRNA with auxiliary pegRNAs that specify triplet payloads (Figure 2a-c). AC repeats were replaced with CCA at the design stage, using its reverse complement, TGG, to provide an opposite-strand NGG PAM for aux targeting while retaining the terminal GGG PAM used by stpegRNA (Figure 2b). The auxiliary primer-binding site anneals to the nicked DNA end, and its reversetranscription template directs payload synthesis [22]. This design couples specified writing to stpegRNAmediated extension and terminal renewal in a write–renew–write architecture. Three aux-payload pools were co-transfected with PE in HEK293T cells on days 0, 7 and 14, respectively, followed by harvest on day 21 for endpoint readout (Figure 2d).

**Figure 2.**
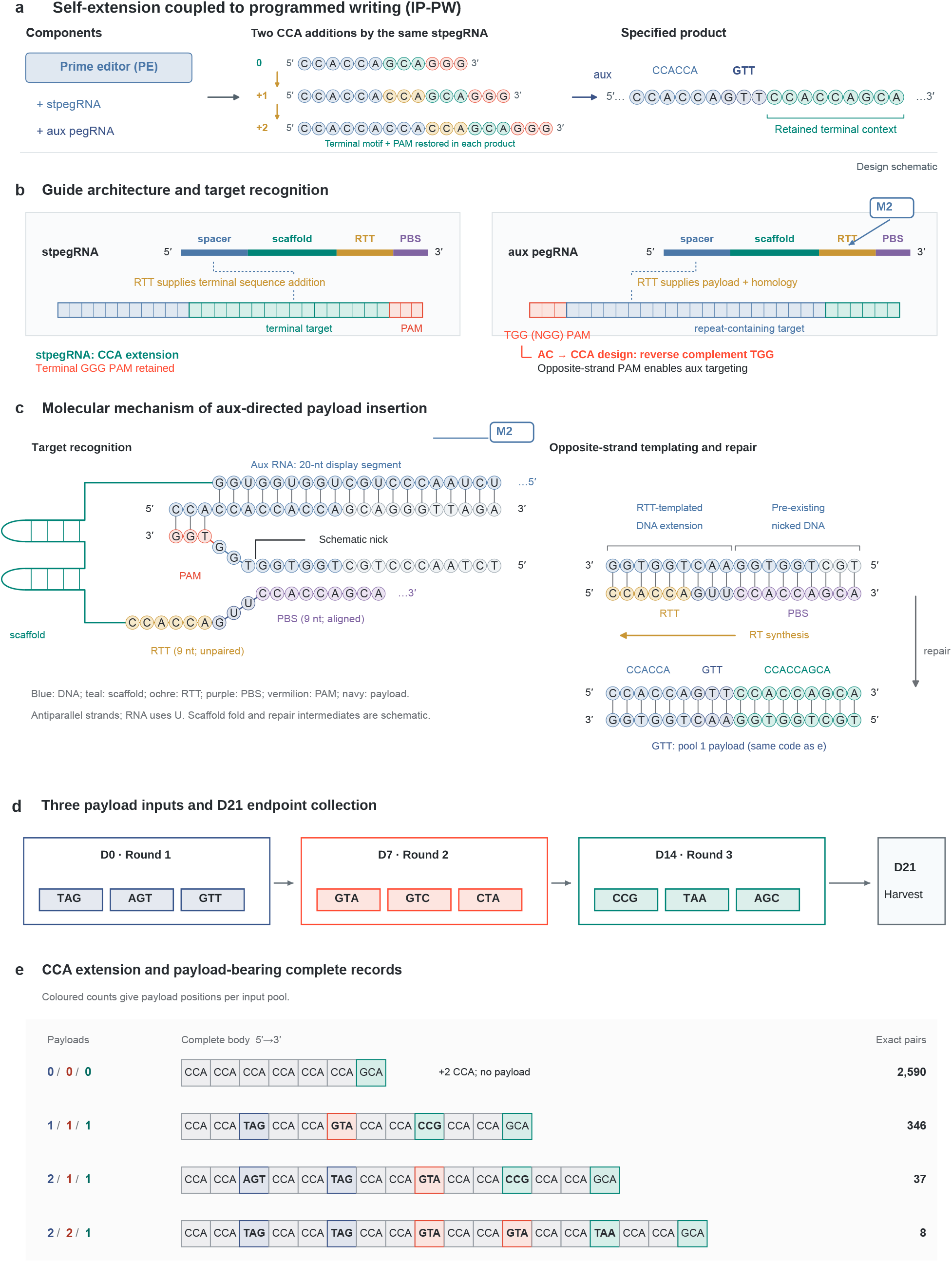
Self-extension coupled to programmed writing. a, The BE-free IP-PW configuration starts with four CCA units and uses transient PE in CCA-stpegRNA-bearing HEK293T cells. b, CCA reverse-complement TGG supplies an opposite-strand aux PAM, distinct from the terminal stpegRNA PAM. c, Aux recognition and templating model with 9-nt PBS, 9-nt RTT and GTT payload example. Nicks and intermediates are schematic. Blue denotes target/spacer, teal scaffold/terminal context, ochre RTT, purple PBS, vermilion PAM and navy the first-pool GTT payload. d, Aux pools introduced with PE at D0, D7 and D14; harvest at D21. e, Complete endpoints with two additional CCA units without payload, or three, four and five payload positions, supported by 2,590, 346, 37 and 8 read pairs. Grey boxes denote CCA and teal outlines GCA. Payload colours identify supplied pools, not measured incorporation times. Counts are PCR-weighted pairs from one biological sample, not independent molecules. Complete distributions are in Figure S2. Matching M2 labels link the aux guide in b to its expanded targeting schematic in c.

Across the complete IP-PW catalogue, we recovered 4,436 distinct, exactly and uniquely parsed sequences containing payload identities from all three input pools, supported by 11,529 read pairs (Figure S2f). Representative complete records contained three, four or five payload positions, supported by 346, 37 and 8 read pairs, respectively (Figure 2e). Payload-free records also extended beyond the four-CCA reference by one, two or three CCA units, with 193,313, 2,590 and 27 mate-concordant read pairs, respectively (Figure S2d); the two-unit gain is also shown in Figure 2e. Each supplied aux template specified one payload together with two CCA units in cassette-forward orientation. None of the displayed three-, four- or five-payload bodies was pre-encoded in a single supplied RTT or complete aux vector. Their distinct payload segments, intervening CCA repeats and retained terminal GCA link accumulated specified content to outputs compatible with terminal renewal (Figure 2c,e and Figure S1f).

Payload-bearing complete records contained one to seven payload positions and accounted for 377,733 of 1,884,163 exactly and uniquely parsed read pairs (20.0%; Figure S2a,e). The 6,283 read pairs containing seven payload positions represented 5,844 distinct complete sequences. Within this set, payload identities from the initial input pool occupied 98.6% of first positions, whereas the first, second and third pools accounted for 49.6%, 35.1% and 15.2% of seventh positions, respectively (Figure S2c). Input-pool labels identify the supplied payload sequences. Together, the multi-payload records, CCA-repeat extensions and retained terminal sequence demonstrate the accumulation of multiple specified payloads within an extending CCA register, supporting the write–renew–write architecture of Infinite Printer. Aux templates also contribute CCA sequence and the initial register contains multiple PAM contexts; endpoint composition therefore supports the architecture without uniquely resolving the sequence of regeneration events.

### Extension retains upstream marks and adds endpoint distinctions

Five complete records of 19, 21, 23, 25 and 29 bp retained a shared 5*^′^*-(AT)3 mark and terminal GCA while differing in intervening AC content (Figure 3b). Each occurred in transcriptome-derived cDNA, non-UMI DNA and UMI-tagged DNA (Figure S3f). Across eight non-UMI libraries harvested on day 7, first-eight-unit marks were 1.72to 7.23-fold more frequent among expanded than reference-length records (Figure 3c).

**Figure 3.**
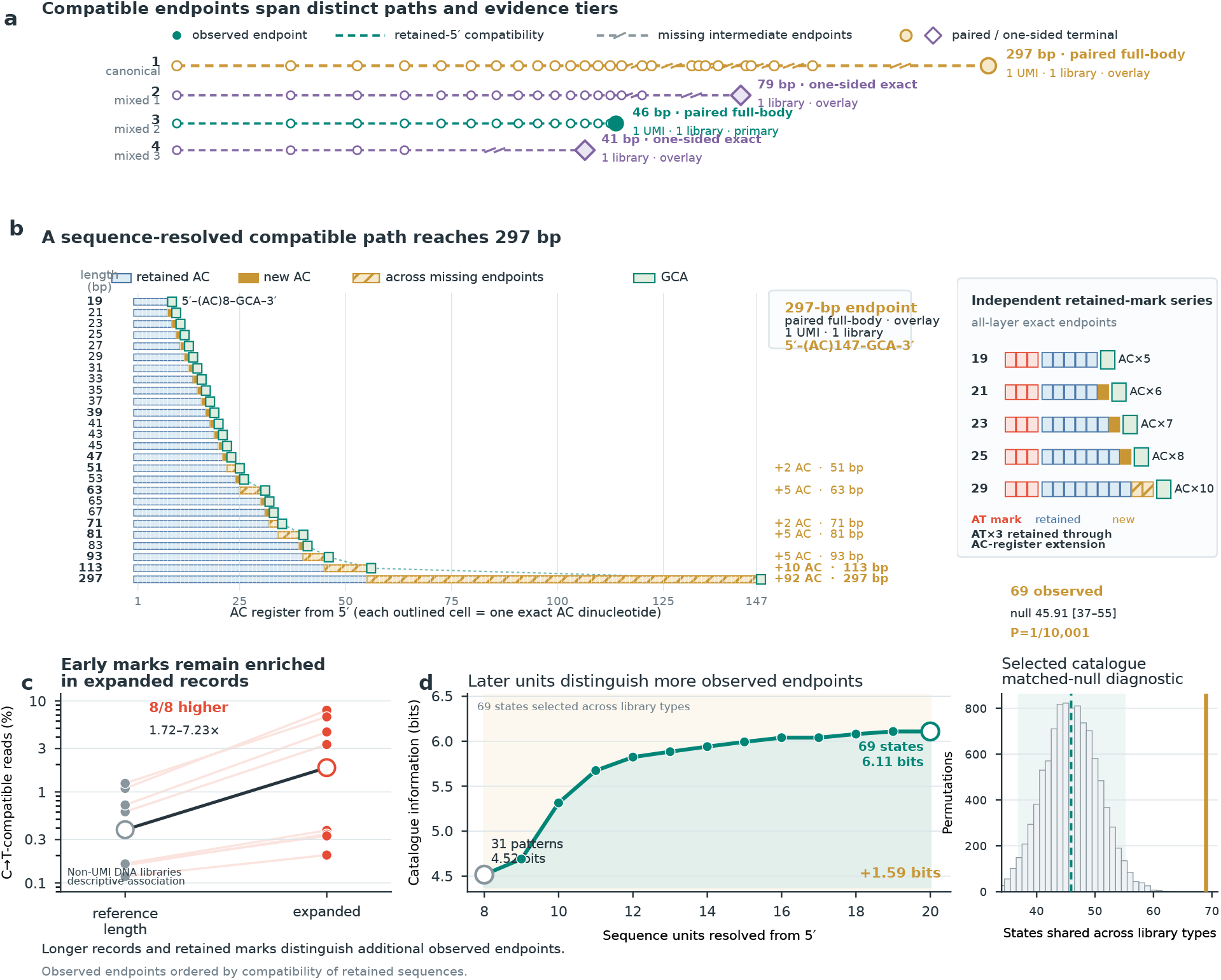
Compatible extension retains marks and adds endpoint resolution. a, Four paths of observed, retained-sequence-compatible endpoints. Dashed connections and explicit gaps represent missing intermediates; terminal labels retain support tier and graph membership. b, Left, 26 endpoints from 19 to 297 bp. Blue denotes retained AC, ochre newly resolved sequence, hatching unobserved intermediate sequence and teal terminal GCA. The 297-bp endpoint has one paired full-body UMI candidate in one library. Right, five endpoints retaining a 5*^′^*-(AT)3 mark across 19–29 bp, each recovered in cDNA, non-UMI DNA and UMI-tagged DNA. c, First-eight-unit marked-read fractions within reference-length and expanded designed-register classes in eight dishes harvested on day 7: four BE-only and four PE+BE. Lines join measurements within a library; large symbols are medians. Measurements summarize within-library marked-read fractions. The separate unpaired dish-level tests are reported in Results and Source Data. d, Equal-endpoint prefix information for 69 states detected in all three library types. The accompanying selection-conditioned overlap diagnostic has observed 69, null mean 45.91, 95% range 37–55 and upper-tail P=1/10,001. The null comparison is conditional on catalogue selection. Paths represent sequence compatibility; ancestry requires independent validation. See Figures S3 and S4.

Four BE-only and four PE+BE dishes started from (AC)8GCA. At complete-body Qeff*≥*20, every PE+BE dish exceeded every BE-only dish for both extension and marked-extension fractions. With all eligible complete-body occurrences as denominator, median compatible extension increased from 5.82% to 6.58%, and extension bearing at least one first-eight-unit C-to-T-compatible mark increased from 0.015% to 0.369%. Mean differences were 0.823 and 0.353 percentage points; exhaustive two-sided dish-label tests gave P=0.0286 each and Holm-adjusted P=0.0571. Median and 95th-percentile body lengths were 19 and 21 nt in both groups; median dish-level 99th percentiles were 21 versus 23 nt (Source Data). These are configuration-level comparisons at the actual editor doses.

Longer records also occurred beyond this series. Library 1 contained 11 canonical states of 31–35 nt with 43 cell–state links; Library 2 contained nine of 31–55 nt at bead level. Library-wide HQ-family support was 93 and 39, respectively. Non-UMI and UMI-tagged DNA catalogues retained longer states and their assay-specific support. Four displayed compatible paths reached 297, 79, 46 and 41 bp, retaining gaps for missing intermediates (Figure 3a; Figure S3). The 297-bp endpoint had one paired full-body UMI candidate in one library; the other termini also had singleton support, with paired or one-sided evidence reported separately. It is the longest observed endpoint, not the sole evidence for extension.

After comparing published editor studies, we selected NG-ABE8e and AncBE4max for experimental evaluation [23, 53]. Symmetric A-to-G/C-to-T rules evaluated complete configurations against AC8 (Figure S4a–c). In the full-body Q20 substitution-compatible subset, AncBE4max showed C-to-T-compatible differences in 90,750/389,281 occurrences (23.31%), including 56,423 with multiple differences; 386 had A-toG-compatible differences. NG-ABE8e showed A-to-G-compatible differences in 9,478/52,623 occurrences (18.01%), including 9,336 with multiple differences and 6,180 with both conversion types. Figure S4a retains Other in its all-letter denominators of 393,868 and 53,089. These layers are distinct; within the compatible subset, each target event includes its exclusive class plus both.

Target-marked AC-compatible bodies longer than 19 nt comprised 12,429 C-to-T-marked occurrences and 46 sequences for AncBE4max, and 1,868 A-to-G-marked occurrences and 29 sequences for NG-ABE8e (Figure S4b). These sequence-stratified counts describe PCR-weighted read support. NG-ABE8e supports an alternative character alphabet, whereas AncBE4max matches the AC/AT representation. No experimental samples or sequencing libraries were excluded from this comparison; analytical eligibility remained sequence based.

At a common depth of 53,089 all-letter, 19-nt, full-body Q20 occurrences, NG-ABE8e had 107 strings and an effective Shannon state number of 3.940. Its technical distribution was a point because this depth used its entire stratum. Across 200 draws without replacement, AncBE4max yielded mean richness 72.3 (technical 2.5–97.5% range 65–78) and effective states 2.920 (Figure S4c). Each configuration contributed one non-UMI library. Thus NG-ABE8e had higher observed diversity in this limited comparison. We selected AncBE4max for subsequent characterization and joint readout because its predominantly C-to-T-directed output maps directly to AC=0 and AT=1, separating length, pattern and retention properties. Broader alphabets provide compatible alternatives for other recording tasks.

Within 69 expanded endpoints detected in all three library types, the first eight units resolved 31 patterns and 4.52 bits of equal-endpoint Shannon information. Reading up to 20 units resolved all 69 endpoints and 6.11 bits, adding 1.59 bits (Figure 3d). Downstream sequence therefore distinguishes records sharing an initial prefix.

Sampling affected compatible-path continuity. Among 116 exact states, a strict one-unit predecessor rule recovered 12, eight and two marked families spanning at least two, three and four nominal steps, but no complete five-step family. Allowing one missing intermediate recovered five steps (Figure S4d,e). In the separate discovery cohort, 4,166/6,137 cell–state links had one observed UMI and 2,482/4,157 cells contained only single-UMI states. Support thinning and cell subsampling reduced family recovery (Figure S4f,g). These sets differ from the final 12,302-cell cohort. Missing intermediate observations therefore do not directly establish cessation of writing.

### Complete records define cell-linked combinations

One stpegRNA design generated 1,549 exact records forming 7,669 combinations across 12,302 transcriptome-linked Library 1 cell labels (Figure 4b,c). An independent bead-to-cell translation linked 20,554 HQ beads to 12,302/13,743 expression-cell labels (89.5%; Figure 4a). Relative to each cell’s largest contributing bead set, aggregation added records in 4,491 cells (36.5%) and 11,933 cell–record links (Figure S5a–c). All observed combinations and their member cells are retained.

**Figure 4.**
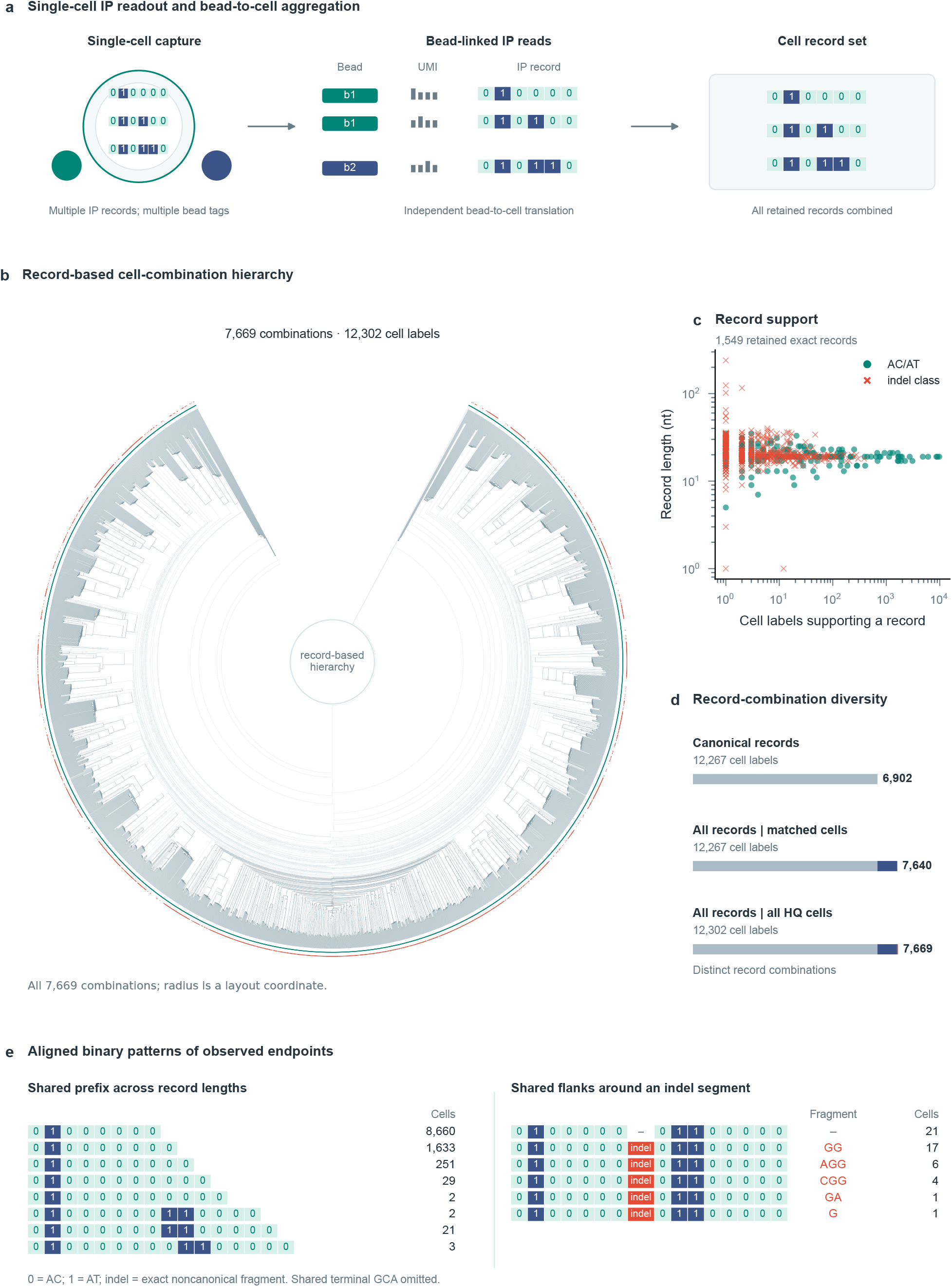
Multiple records define cell-linked combinations. Library 1 observations are independently translated expression-cell labels; counts are descriptive. a, Aggregation of all qualifying records across beads assigned to one cell. b, Complete hierarchy of 7,669 combinations across 12,302 labels, including all 15,336 edges. Joins distinguish shared complete records, shared features and unresolved connections. Teal tips mark combinations; vermilion marks indel-containing combinations. Radius is a layout coordinate. c, All 1,549 records by length and distinct-cell support on logarithmic axes. Circles denote canonical records and crosses indel-class records. d, In 12,267 canonical-positive cells, additional records increase combinations from 6,902 to 7,640; 35 indel-only cells add 29 combinations. e, Eight canonical records with a shared eight-unit prefix and a comparator aligned to five noncanonical fragments. AC=0 and AT=1; indel-labelled cells retain exact strings. Dashes are alignment placeholders and trailing spaces lie beyond observed endpoints. Common terminal GCA is omitted visually. Supporting cohorts, independent Library 2 and molecular matches are in Figure S5.

Canonical records distinguished 6,902 combinations among 12,267 cell labels. Adding exact indel-class records increased this to 7,640 within the same cells, adding 738 distinctions. Another 35 indel-only labels contributed 29 combinations, yielding 7,669 across 12,302 labels (Figure 4d). Library 2 was an independent biological replicate under the same conditions and also underwent single-cell capture, but only recorder-containing cDNA was sequenced. Without a paired transcriptome, its units were capture beads. It recovered 1,174 records and 5,374 combinations across 31,502 HQ beads (Figure S5f–h), independently demonstrating record and combination diversity at the bead level.

Binary matrices further illustrate how complete records distinguish endpoints sharing local sequence content (Figure 4e). Eight canonical AC/AT records shared their first eight units yet retained distinct complete sequences through differences in downstream patterns and record length. A second alignment compared a canonical record with five records sharing the same flanks but carrying different noncanonical fragments. Only interpretable AC and AT units are displayed as 0 and 1, respectively; noncanonical fragments are labelled indel for display while retaining their exact nucleotide sequences. Support for each record was counted as the number of distinct expression-cell labels carrying that sequence. Three exact matches of C4 tag, UMI and complete recorder sequence passed the existing quality criteria in both short-read and ONT observations (Figure S5i). These observations make the additional endpoint distinctions carried by complete sequences directly inspectable and provide sequence-level correspondence across readout platforms.

To examine how complete records support candidate relationships between cells, we organized the 7,669 observed Library 1 record combinations into a hierarchy based on shared exact records and sequence features (Figure 4b). Feature prevalence was evaluated across the cohort of 12,302 cell labels, while undetected records incurred no penalty. Candidate-combination scoring identified a single highest-scoring alternative for 1,379 cell labels; 10,840 had tied highest-scoring alternatives supported by positive shared evidence, and 83 lacked shared variable evidence with another combination (Figure S5d). This analysis identified preferentially supported candidate relationships while retaining multiple alternatives compatible with the observed records. We further assessed sensitivity to within-family state multiplicity by requiring each retained cell–record link to be supported by at least one bead–UMI family containing a single HQ state. This restriction retained 11,910 cell labels and 6,427 record combinations (Figure S5d). Families containing multiple HQ states were reported separately without replacing the primary calls. Together, these analyses organize cell-level combinations through shared complete-record evidence while preserving unresolved candidate relationships.

### Joint spatial recovery of complete molecular records

Recorder-bearing 4T1 cells were implanted into the right first mammary fat pad of a BALB/c mouse. One tumour section underwent spatial profiling and two technical ONT preparations of recorder-containing cDNA, linking records to coordinates and RNA programmes.

The scaffold-upstream spacer defined the primary record; PBS/RTT regions were catalogued separately (Figure 5a; Figure S6a,b). Broad extraction yielded 2,944 spacer strings, including 1,578 linked to coordinates. Recovered UMI, body Qeff*≥*20 and one unambiguous spacer per pooled spot–UMI key defined 297 core states, 15,352 keys and 11,330 spots (Figure 5c,f; Figure S6c,e–g). Among 17,107 eligible keys, 1,755 competing-state keys remained unresolved. Multiple records per spot were retained as sets, defining 842 combinations in a complete hierarchy (Figure 5b; Figure S7).

**Figure 5.**
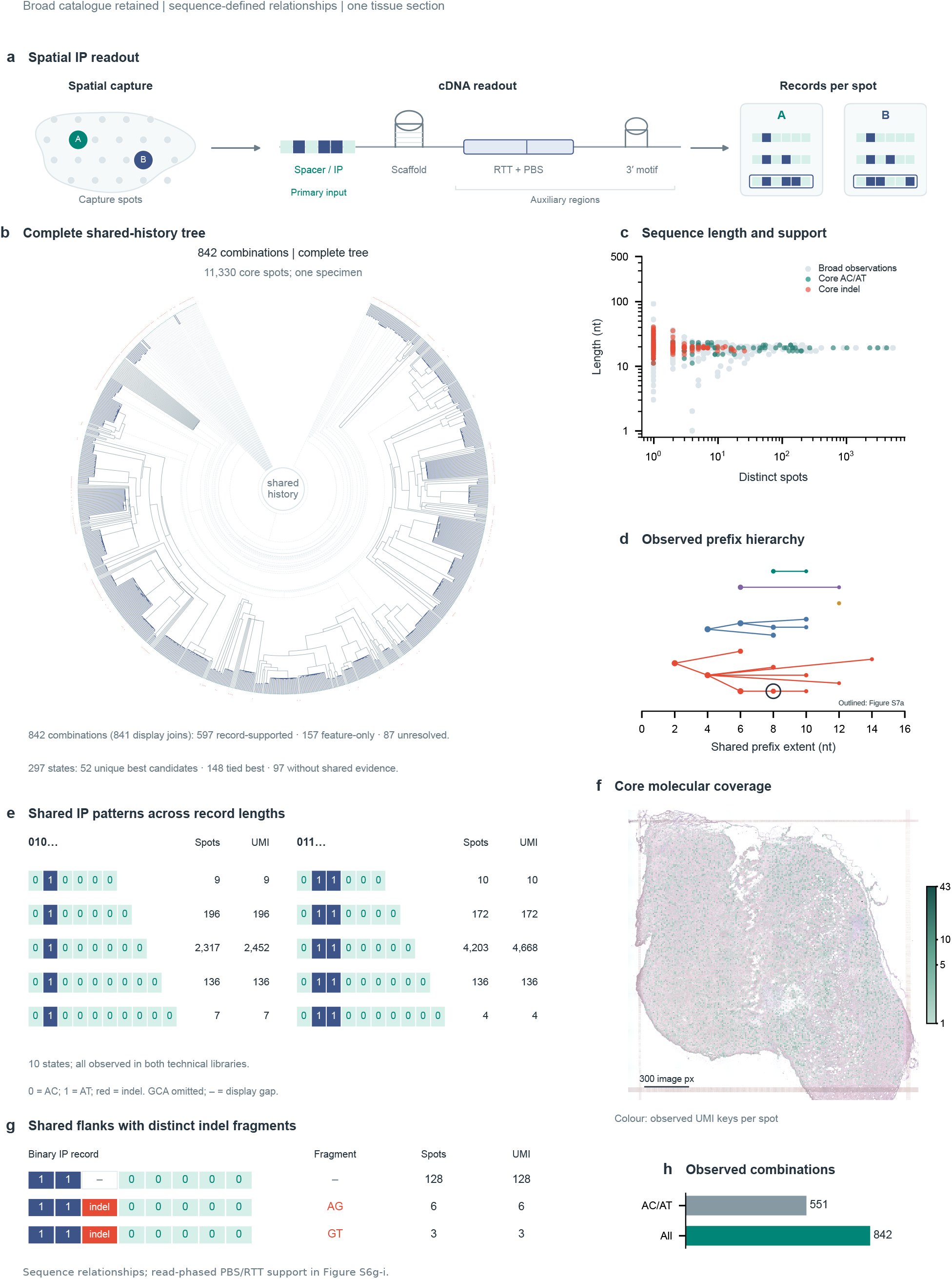
Joint recorder and spatial-transcriptomic readout. One tumour section was assayed using two technical ONT libraries. a, Polyadenylated recorder-containing transcripts support spatial capture and targeted cDNA enrichment; downstream PBS/RTT variants remain auxiliary evidence. b, Complete hierarchy of 842 combinations across 11,330 spots, with 597 shared-record, 157 shared-feature and 87 unresolved joins. c, Length and distinct-spot support for 1,578 located broad strings and 297 core states. Core eligibility requires Qeff*≥*20, recovered UMI and one state per pooled key. d, All 21 variable-prefix histories; node size denotes spot support. e, Ten 010/011-prefix records spanning six to ten units, all detected in both libraries. Support units are spots and pooled spot–UMI keys. f, Molecular support at measured centres over H&E; 15,352 keys across 11,330 core spots, with all 28,739 tissue spots retained as context. Hexagons are display glyphs; ruler, 300 image pixels. g, Canonical comparator and AG/GT variants at 128/6/3 spots; binary and gap conventions follow Figure 4e. h, Canonical-only and all-core combination counts, 551 and 842. Radius in b is a layout coordinate. See Figures S6–S8.

All 841 hierarchy joins were annotated: 597 had informative complete records shared by every descendant, 157 had other shared variable features and 87 had no universal informative witness (Figure S6d). Among 297 states, 52 had one highest-scoring candidate, 148 had ties and 97 lacked shared evidence. Separately, 21 variable-prefix histories described shared 5*^′^* content (Figure 5d). These views organize molecular evidence while retaining unresolved relationships.

Two canonical series sharing 010 or 011 prefixes spanned six to ten units; all ten exact states occurred in both technical libraries (Figure 5e; Figure S7e). Shared-flank noncanonical variants retained AG and GT fragments (Figure 5g), with GC and AA variants in Figure S7d. All core records distinguished 842 spot-level combinations versus 551 for regular AC/AT+GCA records alone (Figure 5h), showing complementary distinctions from length and exact noncanonical content.

Of 15,550 core spacer read observations, 13,759 matched the regional inventory by physical read identity; 8,824 PBS and 9,832 RTT calls met per-base Q30 (Figure S6g–i). Repeated regional variants ranked existing ties for 26 spacer states without changing core calls or candidates. No edge retained support under the diagnostic *≤*1% variant-frequency rule. Corresponding PBS/RTT changes were absent from separate HEK293T DNA datasets supporting Figure 3, which were not matched DNA from this tumour. These variants remain auxiliary cDNA evidence, not established heritable DNA events.

### Information and candidate relationships in complete sequences

To separate information carried by complete records from assumptions about their formation, we analysed the spatial catalogue using a sequence-preserving binary representation. Reliable AC/AT units were represented as 0 and 1, while exact sequences, stable identifiers, register length, terminal status, structural features and ambiguous positions were retained separately. Of 297 core records, 102 were canonical and 81 additional records met the matched-model criteria; the remaining 114 had uncertain register length or terminal status and were retained in the catalogue but excluded from that comparison (Figure S8a–c).

We first quantified the endpoint distinctions provided by different representations of the same 102 canonical records. A visible eight-unit prefix, retaining observed boundaries for shorter records, distinguished 69 classes. Explicitly restoring full register length increased this to 100 classes, while complete sequences distinguished all 102 records (Figure S8f). With equal endpoint weights, the prefix contained 5.902 bits of information, compared with 6.672 bits for complete records. Full length contributed an additional 0.732 bits, and distal character patterns conditional on prefix and length contributed 0.039 bits, giving a total gain of 0.771 bits. Under pooled-UMI weighting, complete records contained 3.294 bits, comprising 3.066 bits in the prefix, an additional 0.227 bits from length and 0.001 bits from distal patterns, for a total gain of 0.228 bits (Figure S8g).

At a fixed register length of eight units, we also recovered 43 distinct character patterns. Conditional pattern entropy across lengths was 4.174 bits with equal endpoint weights and 2.677 bits with pooled-UMI weights (Figure S8e). In a separate matched cohort of 73 records with supported structural information, restoring reliable exact structural identity increased distinguishable classes from 67 to 73, adding 0.164 and 0.023 bits under the respective weighting schemes (Figure S8h). Thus, record growth, distal character content and supported structural identity provided distinct, measurable contributions to endpoint discrimination, with length accounting for most of the information gained beyond the visible prefix in the canonical spatial catalogue.

Independent-character and ordered-extension models used the same 183 endpoints, evidence, scores and candidate pool. Both retained C-to-T directionality; ordering added noncontraction and an exploratory eight-unit terminal-window rule (Figure S9a,b). Of 12,460 chemistry-compatible directed pairs, noncontraction excluded 3,576 and the terminal-window rule another 136, leaving 8,748. Unique-best endpoints decreased from 51 to 49. Full graphs and matrices retained ties, cycles and unplaced endpoints (Figure S9c–f): stronger formation constraints narrowed candidates without increasing unique resolution.

Masking 20% of known positions gave median candidate-set Jaccard similarities of 0.479 and 0.502 for the independent and ordered models. Losing 40% of pooled keys reduced state coverage to 0.781 despite increased conditional stability among survivors (Figure S9g,h). In simulations with 20 genealogies per scenario, ideal terminal editing increased median near-candidate precision from 0.271 to 0.417 with ordering. A wider true editing window instead caused the ordered model to exclude 19.0% of true sampled-ancestor relations. Even under ideal generation, its median wrong-relation fraction among unique-best endpoints was 0.475 (Figure S9i–l). Model agreement with generation and observation completeness therefore affect candidate reliability.

### Spatial and transcriptomic indexing of sequence-defined records

Records were mapped at measured coordinates with all 28,739 tissue spots as context (Figure 5f). An eight-state 1111-prefix group occupied 137 spots (Figure 6a,b). Six of seven first-ring edges joined identical states and one joined different states. Its nominal P=0.0258 did not survive correction (BH q=0.9804; Figure S7a,c). A separate leading-G record occurred at 13 spots, each with one UMI (Figure 6c; Figure S7b).

**Figure 6.**
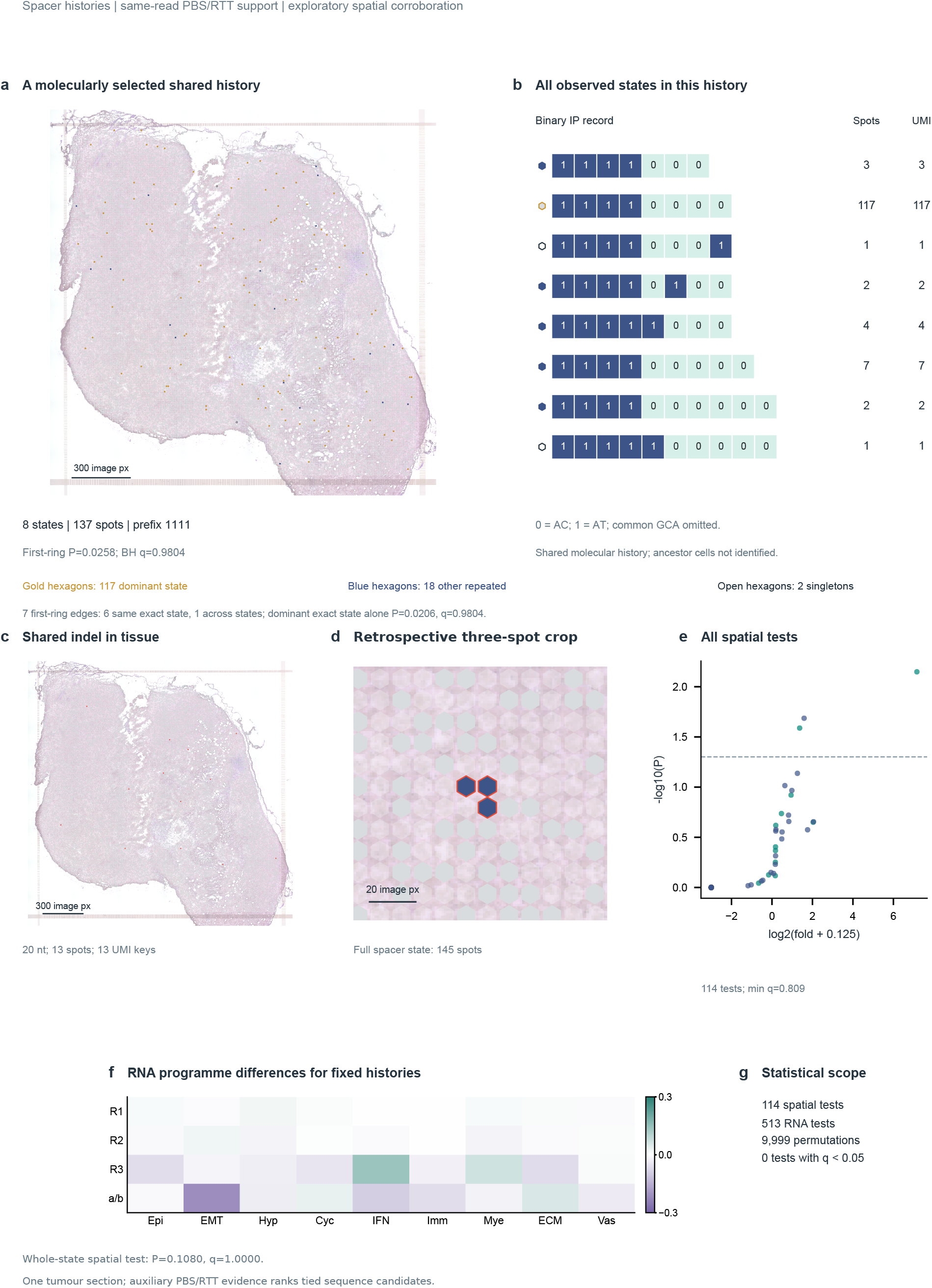
Molecular records across tissue and RNA space. a,b, Eight-state 1111-prefix group at measured centres and as complete binary records, with terminal GCA omitted and no padding beyond endpoints. Gold denotes the dominant state, navy other repeated-UMI states and open glyphs singletons. Group q=0.9804. c, All 13 spots carrying the leading-G record, each with one UMI. d, Retrospectively selected threespot crop; navy marks the complete state at 145 spots overall and vermilion outlines the three locations. Whole-state q=1.0000; ruler, 20 image pixels. e, All 114 spatial tests: log2(observed/null-mean edges+0.125) versus -log10(P). The line marks nominal P=0.05; minimum BH q=0.8094. f, Conditional programme-score differences for three prefix groups and the a/b example, on common ±0.3 standardized-unit limits. R1/R2/R3 are ACAT/AT/ACACAT. All 513 tests are retained; minimum BH q=0.43092. g, Statistical scope: one section, two technical libraries and retrospective analysis. No association survived its correction family. Display glyphs retain measured coordinates and background spots without smoothing. Complete effects, null ranges, P/q and paired-model tests are in Figures S7–S10 and Source Data.

None of 114 spatial-neighbourhood tests survived correction (minimum BH q=0.8094; Figure 6e; Figure S10a). The state in a retrospectively selected three-spot crop occurred at 145 spots overall, giving P=0.1080 and q=1.0000 for its complete membership (Figure 6d; Figure S10d). None of 513 tests across 57 groups and nine RNA programmes survived correction (minimum BH q=0.43092; Figure 6f,g; Figure S10b). Three-, five- and eight-unit prefixes supplied nested spatial views with overlapping memberships (Figures S7f–h and S8d).

Across 18,165 eligible edges and 9,999 paired whole-spot permutations, the ordered-minus-independent excess over the null mean was +0.040 percentage points for record presence (P=0.4337, BH q=0.6542; Figure S10c). Equal-record and log(1+UMI) weighting likewise showed no corrected advantage. Joint readout thus enabled multiresolution indexing and statistical assessment, without detecting corrected spatial or RNA associations or an ordered-model advantage in this section.

## Discussion

Infinite Printer couples self-extension of a molecular record to regeneration of a writable terminal context, allowing new sequence to be added while earlier record content remains upstream of the renewed terminus. In the stochastic-writing configuration, a repeat-complementary stpegRNA directs PE-mediated extension, while AncBE4max diversifies accessible AC-repeat units through C-to-T-directed changes. In a separate BE-free programmed-writing configuration, a CCA-based stpegRNA and auxiliary pegRNAs combine extension with specified payload insertion. CCA-only expansion, complete records carrying multiple payloads and AC/AT records of different lengths retaining shared 5*^′^* marks provide complementary sequence evidence for record growth, diversification and retention of earlier content. By separating renewal of the writable substrate from character generation, this architecture extends recording beyond the positions available in the initial design.

The character-generation module can also be matched to the desired record alphabet. A-to-G-compatible characters and marked longer records recovered in the NG-ABE8e configuration support the possibility of broader alphabets, and this library showed greater observed sequence diversity in the restricted depth-matched comparison. For the present study, we prioritized the predominantly C-to-T-directed output of AncBE4max, which maps canonical repeat units directly onto the AC=0 and AT=1 representation. This simpler alphabet enabled us to examine the distinct contributions of record length, character patterns and upstream-mark retention. Editor configuration can therefore be selected according to the recording task, while self-extension and terminal renewal remain the common architectural foundation.

This architecture complements DNA Typewriter’s designed tandem substrate, peCHYRON’s propagating insertions and PEtracer’s distributed, kinetically tuned cassettes and independent static-barcode benchmarking [15, 19, 20]. A self-targeting guide renews a compatible terminal context as sequence is added, moving the writable frontier beyond the starting positions. Extension potential offers a prospective route to sustained recording, contingent on appropriate kinetics, persistence and molecular recovery. Regulated guide expression, as illustrated by ENGRAM [24], suggests a way to link signals to aux-directed writing. Developmental lineage applications will require calibration against independently tracked pedigrees.

The informational value of complete records arises not only from increased sequence length and character diversity, but also from the coupling of self-extension to relative preservation of previously written information. In the Infinite Printer design, PE-mediated extension regenerates the 3*^′^*terminal target, progressively advancing the accessible BE window relative to the existing record. As the record grows, earlier C-to-T-marked units become positioned farther upstream from the terminal writing region, providing a structural basis for their relative preservation as a readable 5*^′^* history. The shared upstream AT marks recovered across records of different lengths are consistent with this design. Length growth therefore serves two connected functions: it favours separation of previously written characters from the current writing region while adding readable sequence that further distinguishes records sharing an initial prefix.

In the canonical spatial catalogue, the visible eight-unit prefix accounted for 88.4% of the equal-endpoint information, indicating substantial diversity in earlier character patterns. Among records sharing that prefix, length contributed most of the additional information, with distal character patterns and supported structural identity providing complementary distinctions. These contributions are not simply independent: base editing generates record characters, whereas self-extension advances the writable terminus, progressively positioning earlier content upstream while introducing new distinguishable positions. Thus, the extension potential of Infinite Printer supports not only continued record growth but also continued accumulation of information alongside relative retention of earlier content.

This route to accumulating information through an extensible substrate complements the broad state spaces offered by large base-editing arrays [26]. Theoretical sequence combinations, distinguishable recovered endpoints and validated cell-level lineage resolution nevertheless represent different measures. Estimates from the selected cross-library catalogue and pooled-UMI-weighted analyses depend on their respective cohorts and weightings and should not be extrapolated directly to theoretical recorder capacity. Binary matrices make AC/AT differences readily inspectable, while complete records retain observed boundaries, uncertain positions and exact noncanonical sequences. Coupling continued extension, relative historical retention and complete-sequence readout provides a design foundation for accumulating interpretable records in future large-scale lineage-tracing applications.

Formation mechanisms motivate constraints on candidate histories [27, 28]. Ordering narrowed candidates without increasing unique-best resolution, and simulations showed both gains under compatible generation and loss of true relations under mismatch. Homoplasy, incomplete sampling [30], copy mixtures and uncertain editing windows can leave several histories compatible with an endpoint. Retaining ties, cycles, unplaced records and unsupported joins exposes that uncertainty; perturbation stability describes evidence-supported resolution [29]. Independently tracked pedigrees, as used by intMEMOIR [31], are needed to connect sequence relations to biological ancestry.

Joint readout extends the resource to cellular and tissue context. Independent bead-to-cell translation added records in 4,491 transcriptome-linked labels without forcing a single barcode per cell. In tissue, all records per spot were retained and indexed against RNA programmes and coordinates. Spatial spots may contain material from several cells [32]; these are spot-level observations. No association survived multiplicity correction; spatial profiling was limited to the planned day-28 collection. The present results establish joint readout capabilities for future studies of tissue organization and cellular history.

The complete nucleotide sequences recovered by Infinite Printer provide a directly inspectable representation of recorded information. Within its self-extending repeat architecture, record length, shared 5*^′^* prefixes, positional AC/AT patterns and exact noncanonical segments can be identified and compared across endpoints. Sequence alignments reveal both shared marks and positions of divergence, making the molecular basis of record similarity and mechanism-compatible candidate relationships explicit. These endpoint comparisons do not constitute longitudinal observations of individual molecules, however, and sequence similarity alone cannot establish a unique writing order or biological ancestry.

The long-term goal of Infinite Printer is to record cellular histories across large populations during mammalian development and ultimately resolve lineage relationships. Its central architectural feature is the coupling of sequence extension to renewal of a writable terminal context, allowing earlier record content to be retained while preserving the potential for further writing. Here, programmed CCA payloads and diversified AC/AT records provided complementary molecular outputs, with retained upstream marks and complete-sequence differences that can be directly inspected. Single-cell and spatial-transcriptomic workflows further linked these records to cellular and tissue context.

The progressive extension and local diversification of the recorder spacer also raise the possibility that this architecture could impose a relatively low cytotoxic burden during prolonged recording, although this potential advantage remains untested. Future studies in model organisms will need to evaluate sustained recording activity, DNA-damage responses, cellular viability and proliferation, and possible effects on normal development and lineage composition. Such investigations will help establish the long-term biocompatibility of Infinite Printer and advance its application to mammalian development and large-scale lineage tracing.

### Limitations of the study

Interpretation of the molecular records recovered by Infinite Printer depends on preserving the provenance and support level of each observation. PCR-weighted paired reads, ONT family representatives and pooled spatial spot–UMI keys represent distinct observation units, with different sensitivities to amplification bias, sequencing errors and identifier splitting [33, 34, 35]. Repeat-associated polymerase slippage [36] and the distinction between UMI observations and externally calibrated molecule counts [37] further motivate reporting molecular support within each assay rather than pooling counts across sequencing workflows.

Record growth also introduces the possibility of length-dependent recovery bias. Shorter templates can be preferentially amplified, depending on polymerase, buffer and cycling conditions [51, 52]. In the documented PL110 F2/R3 DNA assay, 19-nt and 297-nt recorder bodies correspond to 651-nt and 929-nt amplicons, respectively, with UMI/adapter tails adding 84 nt. Because this KOD-based assay was not calibrated using length standards, no length correction was applied; longer records may be under-represented, but the magnitude of this effect remains unknown. Polymerase slippage and chimeric products remain possible technical contributors, while stochastic writing and differences in effective cellular editing activity may also contribute to the observed long-tailed distribution. These explanations cannot be distinguished from the available endpoints alone. Mate concordance supports agreement between paired reads but does not exclude PCR-induced mutations or template switching.

Exact cross-platform matches further corroborate recovery of matching molecular keys, although these observations may derive from the same original molecule rather than independent molecules. PBS/RTT variants detected in spatial cDNA were not observed in the separate HEK293T ONT DNA datasets supporting Figure 3, which were not matched DNA from the spatial specimen. Cytosine base editors can act on RNA [38], including reported RNA activity of AncBE4max [39], making RNA editing one possible explanation without establishing the origin of individual variants. We therefore retain PBS/RTT differences as auxiliary cDNA evidence rather than verified heritable DNA edits. The epegRNA motif was included to support guide-RNA integrity [40].

## Resource availability

### Lead contact

Further information and requests for resources should be directed to the lead contact, Demeng Chen.

### Materials availability

Plasmids generated in this study are described in the Key Resources Table and construct resources. Requests for these materials should be directed to the lead contact and are subject to applicable institutional material-transfer requirements.

### Data and code availability

Sequencing collections have been deposited in GSA (CRA051489) and GSA-Human (HRA021111) under BioProject PRJCA074520. Raw sequencing data remain non-public during peer review, with public release planned upon journal publication. The accompanying Source Data contain the processed record tables, analysis cohorts, statistical outputs and figure-specific mappings; Table S1 defines the editor-analysis denominators. Spatial RNA analyses used the original vendor-delivered Infi_1.h5ad. Library 2 has recorder-targeted cDNA data and no paired transcriptome. Confidential access to the deposited data is available to editors and reviewers on request through the editorial office.

Analysis and figure code are provided in Data S2 for confidential peer review and are maintained in the private repository https://github.com/Zhang-Zhihui-Andrew/infi_printer. Public release is planned upon journal publication. The resource index specifies the required processed inputs and their versions. Additional information required to reanalyse the reported data can be requested through the lead contact.

## Acknowledgements

The authors have no additional acknowledgements.

## Author contributions

Zhihui Zhang conceived the study and designed the overall experiments. Zhihui Zhang performed plasmid construction. Zhihui Zhang, Yingyi Li and Yifang Peng performed the cell and animal experiments and prepared sequencing libraries. Kang Li, Yixin Cheng and Maosheng Cheng contributed to the experimental work. Zhihui Zhang analysed the data, with contributions from Xiaochen Wang, and prepared the figures. Zhihui Zhang and Rongsong Ling developed the analysis code. Caihua Zhang contributed to code development. Zhihui Zhang wrote the initial draft. Shuang Chen, Zhenyu He and Demeng Chen reviewed and revised the manuscript. Demeng Chen provided research guidance. Zhenyu He supervised the project. All authors read and approved the manuscript.

## Declaration of generative AI and AI-assisted technologies in the manuscript preparation process

During preparation of this work, OpenAI Codex assisted with drafting analysis and figure scripts, text editing, formatting and file-consistency checks. The authors are responsible for verification of the assisted work and for the scientific content.

## STAR Methods

### Key resources table

The Key Resources Table is provided below. Complete oligonucleotide and construct fields and simulation settings accompany the resource tables.

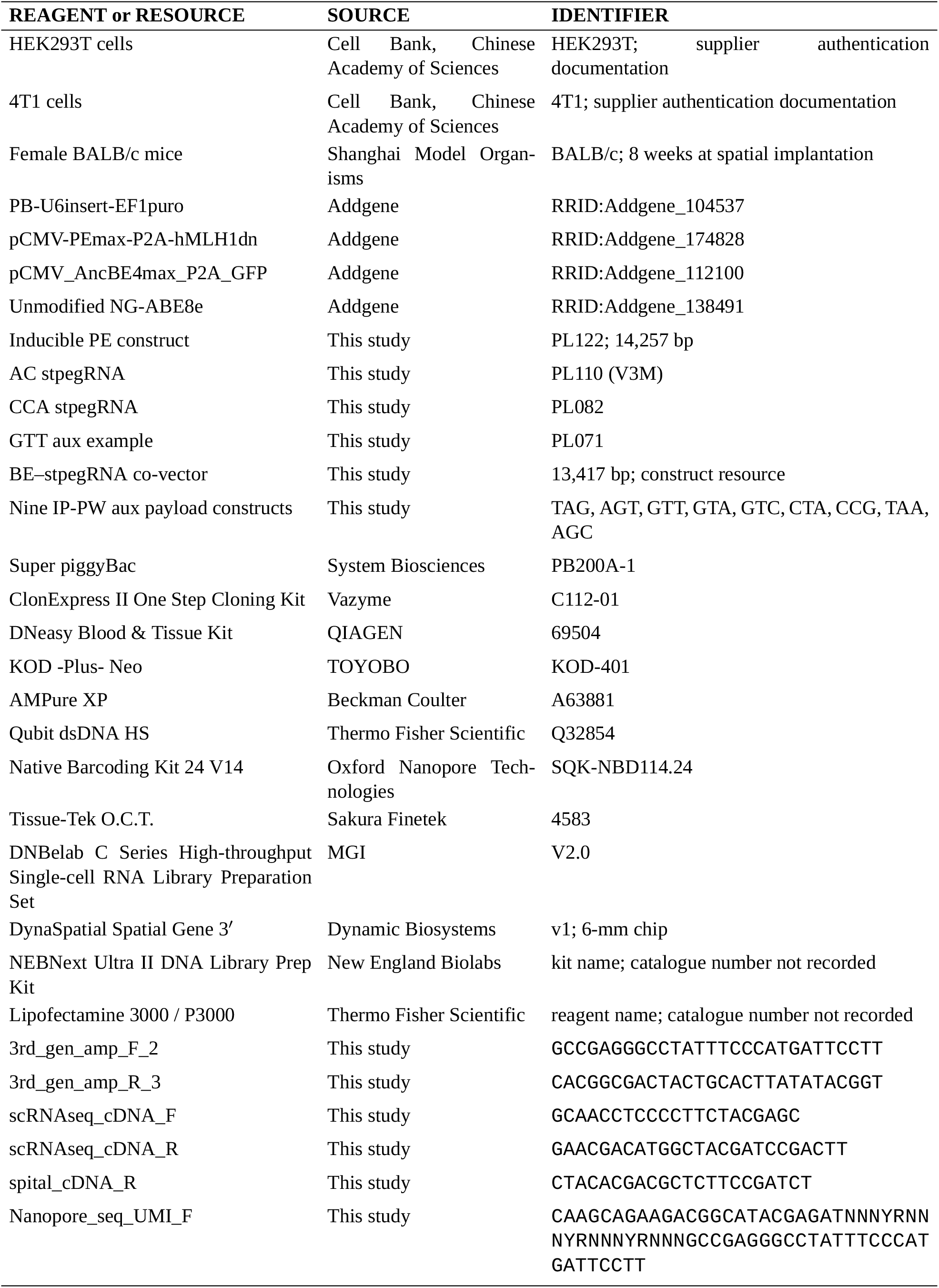
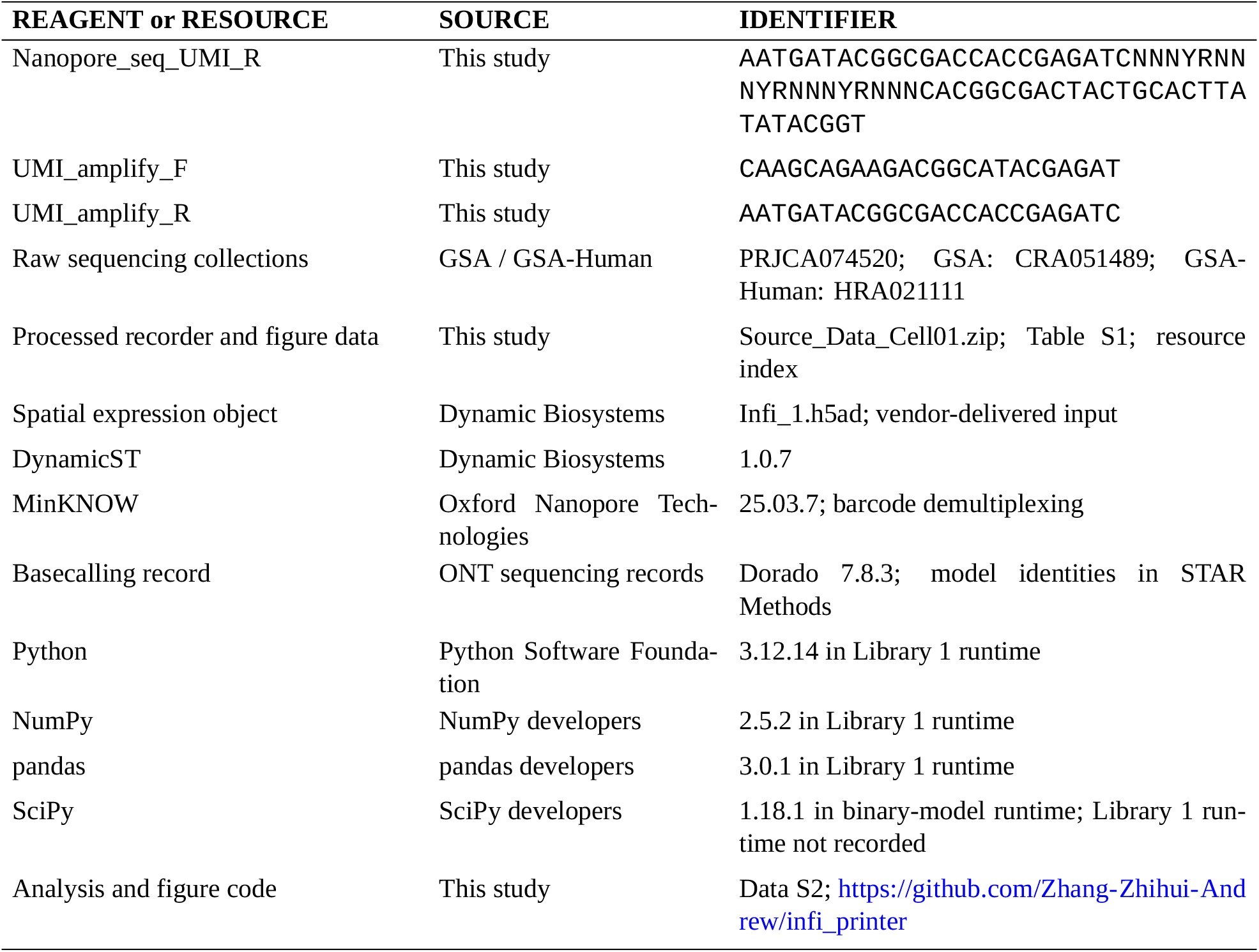

### Experimental model and study participant details Experimental design and observation units

Experiments evaluated self-extension, base diversification, programmed payload writing and joint transcriptomic readout. IP-SW used an AC-repeat recorder with base editing; the separate, BE-free IP-PW configuration used a four-CCA starting register. All plasmid constructs used in this study were verified by Sanger sequencing, and the corresponding sequencing records were retained. The eight-dish extension comparison, editor-configuration comparison, targeted UMI-tagged DNA collection, single-cell experiments and spatial experiment were separate experimental cohorts.

The eight-dish comparison comprised independently cultured HEK293T dishes with BE plus stpegRNA (four dishes; barcode01–04, IP_S001–IP_S004) or PE plus BE plus stpegRNA (four dishes; barcode05–08, IP_S005–IP_S008). Cells stably expressing stpegRNA were distributed before transient editor delivery. The starting recorder body was (AC)8GCA (19 nt), confirmed by Sanger sequencing. Conditions were randomly assigned; the analyst knew group assignments. Cultures were harvested seven days after editor transfection, without post-transfection antibiotic selection. No samples or replicates were excluded from this comparison.

HEK293T Libraries 1 and 2 were independent biological replicates prepared under the same experimental conditions. Both underwent single-cell capture and targeted recorder-cDNA sequencing, but only Library 1 had paired transcriptome sequencing. Library 1 records were aggregated by independently assigned expression-cell labels; Library 2 retained capture-bead barcodes as observation units. The spatial experiment comprised one tumour section from one mouse and two technical ONT libraries. IP-PW used one HEK293T biological sample. No experimental samples or sequencing libraries were excluded from the editor-configuration comparison; read- and sequence-level eligibility criteria were applied as described below.

An exact endpoint is a recovered nucleotide string. A record combination is the unordered set of qualifying distinct strings in a cell label, bead barcode or capture spot; sequence order within each record is preserved. Read pairs, reads, UMI families, bead barcodes, cell labels and spots were counted separately and were not substituted for biological replication. Sample, event and library identifiers link experimental material to sequencing files and panel-level Source Data. Analysis catalogues, including the 69-state, 116-state and 4,157-cell sets, are inclusion cohorts rather than additional biological samples. Expression-sequencing partitions of Library 1 were retained as partitions of one library. Single-cell and spatial recorder libraries were cDNA-derived; the remaining targeted recorder collections used extracted DNA. No biological samples were excluded from the single-cell or spatial experiments.

### Cell culture, transfection and induction

HEK293T and 4T1 cells were obtained from the Cell Bank of the Committee on Type Culture Collection of the Chinese Academy of Sciences, with authentication documentation from the supplier. Both tested negative for mycoplasma and were cultured in high-glucose DMEM containing 10% fetal bovine serum and 1% penicillin– streptomycin at 37 °C and 5% CO_2_.

HEK293T transfection used Lipofectamine 3000/P3000 in Opti-MEM at 70–80% confluence. In the eight-dish comparison, each six-well culture received 2.5 µg total plasmid DNA; PE-containing cultures used a 1:1 molar PE:BE ratio, and neither group received empty-vector balancing. The comparison therefore evaluates the complete editor configurations at these doses. Medium was replaced after 6 h. Cultures were passaged at 80–90% confluence, usually every 2–3 days at 1:3–1:5, without pooling at harvest. Mixing and passaging procedures are described in this subsection.

Stable transcript-linked recorder lines were established sequentially with Super piggyBac (SBI, PB200A-1). Transposase and transposon vectors were delivered at a 1:2.5 molar ratio and 2.5 µg total DNA per six-well culture. PE delivery and blasticidin selection preceded BE–stpegRNA delivery and puromycin selection. Selection began approximately 48 h after transfection and lasted approximately two weeks per round for HEK293T and one week for 4T1. Blasticidin concentrations were 5 and 10 µg/mL, and puromycin concentrations were 2 and 4 µg/mL, respectively. Only puromycin was maintained in the second round. Integrated recorder copy number was not measured by the recovery assay.

Bulk cultures expressing the original CMV-driven editors received no doxycycline (Dox). HEK293T singlecell cultures received 2 µg/mL Dox after stable-line establishment and selection, with sampling after 30 days of induction. Medium was usually replaced daily with Dox replenishment. The separate 4T1 C4 sample was collected from cultured cells on day 30, not from a mouse tumour. Its day-zero reference and Dox exposure duration were not separately recorded. For the spatial experiment, there was no pre-implantation Dox induction. In vivo schedules are specified below.

IP-PW used stably introduced CCA-stpegRNA. PE and three aux plasmids were co-transfected at a 3:1:1:1 molar ratio on days 0, 7 and 14, followed by harvest on day 21. Day 0 denotes the first payload input; plasmid-clearance kinetics and payload incorporation times were not measured. For the UMI-tagged genomic DNA collection, PE and BE plasmids were replenished every seven days and cultures harvested on day 28 after initial editor transfection.

For six-well HEK293T transfection, DNA and 5 µL P3000 were diluted in 125 µL Opti-MEM; 7.5 µL Lipofectamine 3000 was diluted separately in another 125 µL. Dilutions were mixed, incubated for 15 min at room temperature and added dropwise.

For passaging, cells were washed twice with PBS and detached using 0.25% trypsin at 37 °C for approximately 1–2 min. Serum-containing medium quenched trypsin. Cells were centrifuged at 300*×*g for 3 min and resuspended. Excess cells were discarded during routine passaging, and cultures were not pooled at harvest.

### Mouse experiments and tumour sampling

Recorder-bearing 4T1 cells were implanted percutaneously into the right first mammary fat pad of female BALB/c mice, using 1 *×* 10^5^ viable cells in 50 µL PBS without Matrigel or incision. Mice were eight weeks old at implantation. The spatial mouse was supplied by Shanghai Model Organisms and acclimatized for at least seven days. Isoflurane was used at 4–5% for induction and 1–2% for maintenance; no additional inoculation or post-inoculation analgesic was administered.

For spatial profiling, one tumour was collected 28 days after implantation and one section analysed, without selection among multiple specimens. Its two ONT libraries were technical preparations, not biological replicates. Maximum tumour dimension at collection was 7 mm. The mouse received standard mouse chow and Dox-containing water. After implantation, the mouse received Dox at 2 mg/mL in drinking water until collection; in the spatial experiment this began on implantation day 0. Water was replaced every three days, protected from light and prepared without added sugar.

Health was observed daily on days 1–9, at least twice daily on days 10–21, and every two days from day 22 until collection; weight and tumour size were measured every two days. Humane endpoints followed the UCSF IACUC tumour-study guidelines: a tumour dimension of 20 mm in any direction, restricted mobility or access to food or water, body condition score below 2, or laboured respiration. Ulceration endpoints included bleeding, infection, necrosis, discomfort or failure to stabilize within seven treatment days. Spatial tumour collection occurred at the planned day-28 endpoint. Euthanasia used cervical dislocation under deep isoflurane anaesthesia, with death confirmed before collection. Spatial analysis was not blinded.

All animal experiments were approved by the Clinical Research and Laboratory Animal Ethics Committee, The First Affiliated Hospital of Sun Yat-sen University (approval no. [2026]203). The corresponding authors confirm that these experiments were conducted in accordance with applicable guidelines and regulations.

### Method details

#### Plasmid construction and guide design

The stpegRNA cassette was assembled by annealing complementary oligonucleotides and ligating them into BamHI/EcoRI-digested PB-U6insert-EF1puro (Addgene 104537; RRID:Addgene_104537) [5]. Bulk DNA experiments used the original CMV-driven pCMV-PEmax-P2A-hMLH1dn (Addgene 174828; RRID:Addgene_174828) [41] and, where required, pCMV_AncBE4max_P2A_GFP (Addgene 112100; RRID:Addgene_112100) [23], with separate stpegRNA constructs. IP-PW omitted BE and used separate PB-U6insert-EF1puro-derived plasmids for stpegRNA and each aux payload.

For transcript-linked readout, the PEmax–P2A–hMLH1dn fragment was amplified from Addgene 174828 and assembled into an inverse-PCR-linearized XLone-PE2 backbone using ClonExpress II One Step Cloning Kit (Vazyme, C112-01). The inducible construct contained TRE3GS, Tet-On 3G and an EF-1*α* core promoter in the regulator cassette [50], with blasticidin resistance. The AncBE4max–P2A–GFP fragment was assembled with PB-U6insert-EF1puro, followed by insertion of U6–stpegRNA using the same method. DpnI treatment removed PCR template plasmids before assembly. The 13,417-bp BE–stpegRNA vector carried puromycin resistance, with the recorder cassette downstream of the resistance coding region and upstream of the SV40 polyadenylation signal. Polyadenylated Puro transcripts containing the recorder supported capture-barcode-linked cDNA recovery.

The AC design was stpegRNA-AC (PL110; source alias V3M), and IP-PW used (CCA)4GCA (PL082). The AC guide contained an 11-nt RTT and 14-nt PBS; the CCA guide contained an 11-nt RTT and 10-nt PBS. Domain definitions follow prime-editing terminology [22]. The AC RTT specifies ACGCAGGGTTA in DNA orientation, including one additional AC unit. Targeting/cloning fields include cloning-associated bases. Activity from both repeat arms was not independently measured. PL071 supplies the illustrated GTT-payload aux example.

A 37-nt epegRNA stabilizing motif followed a construct-specific 8-nt linker designed with pegLIT [40]: ATTAATTA for the AC construct and AATATTAA for the v21 CCA constructs. Linker and motif are excluded from RTT/PBS lengths. Guide fields are listed in the Construct and oligonucleotide sequence fields resource; complete construct sequences and maps are associated with their construct identifiers. The AC targeting/cloning field was 29 nt and its annotated scaffold field 72 nt; adjoining construct bases were retained separately. The PL071 example contained a 9-nt RTT and 9-nt PBS. These annotations specify construct sequence domains. All nine aux templates specified one triplet payload and two CCA units in recorder-forward orientation. Complete aux sequences and observed payload intervals were compared, and opposite-strand NGG contexts in the starting CCA register were enumerated as potential starting sites; payload parsing allowed all compatible routes.

#### Targeted amplification and sequencing

Genomic DNA was extracted with DNeasy Blood & Tissue Kit (QIAGEN, 69504). Recorder DNA and cDNA were amplified with KOD -Plus- Neo (TOYOBO, KOD-401), purified with AMPure XP beads (Beckman Coulter, A63881; 0.8:1 bead:sample ratio), and quantified with Qubit dsDNA HS (Thermo Fisher Scientific, Q32854). Primer sequences are provided in the Construct and oligonucleotide sequence fields resource; purification procedures are described below.

Non-UMI DNA libraries 01–08 and IP-PW used one 25-cycle PCR with 50 ng DNA in 50 µL and 0.3 µM each F2/R3 primer. Cycling was 94 °C for 2 min, then 98 °C for 10 s, 63 °C for 30 s and 68 °C for 30 s. Single-cell and spatial recorder-cDNA PCR used 150 ng template, the same reaction volume and primer concentrations, and 25 cycles; annealing was 61 °C and 60 °C, respectively, for 30 s, with 68 °C extension for 30 s. Remaining steps followed the same three-step program. Single-cell reactions paired scRNAseq_cDNA_F/R; spatial reactions paired scRNAseq_cDNA_F with spital_cDNA_R.

The ten UMI-tagged genomic DNA libraries used paired 18-nt patterned tags adapted from Karst et al. [42]. Tagging PCR used 50 ng DNA in 50 µL, 94 °C for 2 min, two cycles of 98 °C for 10 s, 55 °C for 15 s and 68 °C for 2 min, then 68 °C for 10 min. All purified tagged product underwent 25-cycle amplification with universal primers using the same thermal steps; both rounds used 0.3 µM each primer. UMI-family representative selection and singleton eligibility are specified in “UMI endpoints, compatible paths and sampling sensitivity” and “Capture-barcode recovery and cell aggregation”.

ONT libraries for Figure 3–6 were prepared in-house with Native Barcoding Kit 24 V14 (SQK-NBD114.24), including end repair, dA-tailing and native-barcode ligation, and sequenced on PromethION R10.4.1 flow cells at default acquisition settings. Main-figure datasets used the SUP v4.3.0 model for R10.4.1/E8.2 at 400 bases/s. The editor-comparison FASTQ headers specified HAC v4.2.0. Software versions are listed in “Statistical reporting, visualization and reproducibility”. Novogene prepared IP-PW PCR products with NEBNext Ultra II DNA Library Prep Kit and sequenced them on NovaSeq 6000, with 250 nt per end.

AMPure XP purification used 0.8 volumes of beads per sample volume and a 5-min room-temperature binding incubation. Beads were washed twice with 80% ethanol, air-dried for approximately 5 min and resuspended in 22 µL 0.1*×* TE. After 2 min, 20 µL eluate was recovered. Purified products were quantified by Qubit dsDNA HS; fragment size was not assessed for the targeted PCR products.

In the PL110 F2/R3 assay, recorder bodies of 19 and 297 nt correspond to amplicons of 651 and 929 nt before UMI/adapter tails, which add 84 nt. The KOD-based assay was not calibrated using length standards and no length-dependent recovery correction was applied.

#### Single-cell and spatial transcriptomic preparation

HEK293T and 4T1 capture, reverse transcription and cDNA amplification followed the DNBelab C Series High-throughput Single-cell RNA Library Preparation Set V2.0 protocol (MGI) [43]. Cells were emulsified with mRNA-capture beads; after lysis and poly(A)-RNA capture, beads were recovered for reverse transcription, generating barcode- and UMI-tagged cDNA. Library 1 delivery records specified paired read lengths of 30/100 nt for the cDNA library and 20/30 nt for the accompanying oligonucleotide library. Library 2 underwent recorder-targeted sequencing only. Recorder-containing Puro transcripts were recovered through poly(A) capture and targeted enrichment.

Spatial libraries used DynaSpatial (Dynamic Biosystems), based on Decoder-seq [44], with a 6-mm chip and Spatial Gene 3*^′^*v1 poly(A)-capture chemistry. Tumours were embedded in Tissue-Tek O.C.T. (Sakura Finetek, 4583) and sectioned at 10 µm. Following the published workflow [32], sections were methanol-fixed for 30 min, H&E-stained, imaged, permeabilized and reverse-transcribed on the array. Released cDNA was preamplified and amplified using a cycle number determined by qPCR. Amplified cDNA was split between expression-library preparation and two recorder PCRs (150 ng each; barcode12/13). Expression-library preparation comprised fragmentation, end repair, A-tailing, adapter ligation and dual-index PCR.

DynamicST 1.0.7 processed expression data against mouse mm10 with intronic counts. The delivered Infi_1 expression object contained 28,739 tissue-covered spots, 32,285 gene features and eight clusters among 36,864 measured array positions. Leiden settings were resolution 0.8 and random state 0 [45]. These labels supplied expression context; vendor Seurat and BayesSpace outputs were not used to calculate the RNA programme scores. Recorder calls were linked to measured coordinates without converting capture spots into individual cells.

### Quantification and statistical analysis

#### Complete-state calling and the eight-dish comparison

Reads were searched in both orientations using design-derived landmarks. Complete intervening bodies were normalized once to recorder-forward orientation, preserving original orientation, source IDs and exact strings. Partial or terminal-only matches did not replace complete bodies. Sequence IDs were linked explicitly to strings; the earlier catalogues used the first 20 hexadecimal characters of SHA-256 digests.

For the eight-dish comparison, all exact leader/scaffold pairs were enumerated and three candidate PAM bases separated. No AT requirement, length cap or selected-state catalogue preceded extraction. Distinct body/PAM interpretations remained ambiguous; identical interpretations were counted once using the least favourable quality. The primary denominator contained all nonempty A/C/G/T bodies with a unique interpretation and Qeff *≥*20, including noncanonical bodies, where *Q*_eff_ = *−*10 log10[mean(10*^−Q^*^/10^)]. Unfiltered and full-body per-base Q20/Q30 layers were sensitivity analyses.

Design-compatible extension required (A[CT])nGCA with n >8. Marked extension additionally required at least one T in the first eight units. Both primary fractions used each dish’s eligible complete-body occurrences as denominator. Extension conditional on canonical bodies and length distributions were reported separately. Complete length inventories retained empirical cumulative distributions, unique-body support and length-bymarking counts, including zero bins, rather than restricting analysis to the displayed endpoint series. All 70 allocations of four PE+BE labels among eight dishes were enumerated; two-sided tests compared the absolute difference in mean dish fractions, with Holm correction across the two primary endpoints. Medians and length percentiles were descriptive. Non-UMI counts remained PCR-weighted read occurrences.

For within-library mark enrichment, the first eight units had A at each A position and C/T at each C position, with at least one T; terminal GCA was excluded from this window. Marked-read fractions were calculated separately among reference-length and expanded designed-register bodies. This descriptive comparison did not treat all eight libraries as replicates of one editor condition.

#### Programmed-writing endpoint analysis

The IP-PW reference was (CCA)4GCA. Payload pools, in recorder-forward orientation, were TAG/AGT/GTT on day 0, GTA/GTC/CTA on day 7 and CCG/TAA/AGC on day 14. All endpoints were collected on day 21. Pool labels identify supplied sequences and input days, not measured incorporation times.

Both read orientations were processed with the exact-flank caller, requiring mate-concordant complete bodies. The 2,220,942 synchronized pairs yielded 1,884,170 concordant occurrences. Design-aware parsing assigned CCA to the register, used the first eligible boundary at zero-based position 6, and retained unique exact parses with zero residual edit distance and no junction-like flag. Ties were not forced to one interpretation; the eligible payload-depth denominator was 1,884,163 pairs.

Pure (CCA)+GCA strings were counted separately. CCA gain was measured relative to four repeats after separating payload positions. Identity-presence summaries counted each identity once per pair and could overlap. Position-specific pool composition used the same 6,283 seven-position pairs at every position. Support was PCR-weighted read-pair occurrence, not independent molecule count. The supplied templates and starting-register PAM contexts were evaluated alongside endpoint composition rather than assuming that endpoint structure uniquely determined the order of regeneration events.

#### UMI endpoints, compatible paths and sampling sensitivity

For Figure 3 UMI-tagged DNA, the first and last 300 nt of each read were searched in both orientations. Inner primers were recovered with a seeded Levenshtein verifier allowing one substitution, insertion or deletion; this tolerance applied to primers, not exact endpoint identity. Adjacent 18-nt tags had to match the patterned grammar. Two valid ends defined paired support; one valid end could define a candidate when the opposite end was absent or damaged.

Body recovery preferred exact dual boundaries, then scaffold/PAM or terminal rescue, followed by endpoint-anchored AC/AT register matching with at most two small substitutions or indels. Recovery tier was retained. No UMI-neighbour clustering, family consensus or singleton exclusion was applied. Endpoint IDs with incompatible states were excluded from the unique-state catalogue but retained for inspection. Endpoint-defined candidates were not assumed to be independent molecules because identifier errors can split templates [35].

Compatible paths preserved ordered 5*^′^* sequence, allowed AC-to-AT changes and terminal extension before GCA, and displayed unobserved intermediate lengths as gaps. Evidence tier and technical-library support remained attached to every endpoint. Links represented sequence compatibility, not tracked transitions. Retained-mark endpoints were matched exactly across transcriptome-derived cDNA, non-UMI DNA and UMI-tagged DNA.

The 69-state information catalogue required detection in all three library types. Endpoints were equally weighted and projected to their first min(u, observed units) units; Shannon information was calculated from projected class frequencies [46]. The 10,000-draw length/category-matched overlap permutation was therefore selection-conditioned, not an independent recurrence test.

A separate 116-state (AC|AT)+GCA catalogue tested incomplete sampling. Strict predecessors removed one terminal AC unit; relaxed searches allowed up to three missing length steps. Marked families were evaluated at depths of two to five steps. The exact-margin null preserved sequence count, length, AT count and positional AT frequencies, using four chains with seed 20260830, burn-in 10,000 and thinning 1,000 for 10,000 retained catalogues. The exploratory grid was not multiplicity-adjusted. Support thinning and cell subsampling used eight retained fractions from 1 to 0.01 and 1,000 repeats, with seeds 20300830 and 20340830, respectively. The latter used a separate discovery cohort of 4,157 cells, 65 states and 6,137 links, not the Figure 4 cohort. Library breadth described technical support within the selected catalogue, not independent biological recurrence. The eight-dish mark comparison and editor-character analysis retained their own experimental inputs and denominators.

#### Editor-configuration comparison and diversity

After comparing published studies of different base editors, we selected NG-ABE8e and AncBE4max for experimental evaluation of adenine- and cytosine-base-editing configurations, respectively [23, 53]. Their A-to-G and C-to-T activities provided a mechanistically informative contrast for diversification of the AC-repeat recorder. AncBE4max and unmodified NG-ABE8e (Addgene 138491; ecTadA(8e)–nSpCas9(NG)) were evaluated with matched stpegRNA, PE configuration and sampling conditions. The comparison concerned complete editor configurations, including their Cas components, and was separate from the eight-dish experiment. AncBE4max was used for binary recording because its C-to-T-directed spectrum matched the AC/AT alphabet.

The complete merged FASTQ files be_6.fastq.gz and tad_6.fastq.gz represented the AncBE4max and NGABE8e analysis inputs, respectively. One non-UMI sequencing library per editor configuration was analysed; merged files and read counts were not treated as biological replicates. Both files were processed in full using the same extraction and quality rules. In each orientation, the caller used the first exact AACACCGG leader and first subsequent TTAGAGCTAGAAATAGCAAGTTAACCT scaffold, separated the preceding three PAM bases, and admitted bodies up to 10,000 nt. Candidate choice favoured GGG PAM, then length closest to 19 nt, then forward orientation. This differed from the eight-dish all-boundary caller.

The reference was complete (AC)8GCA, including terminal positions. Full-body layers required every base to meet Q0, Q20 or Q30; site-specific spectra applied the threshold only at that site and counted all alleles. All-letter analyses retained every eligible 19-nt string. Substitution-compatible bodies allowed only unchanged bases, A-to-G and C-to-T substitutions, without gaps or other substitutions. Joint classes were unchanged, A-to-G only, C-to-T only, both and Other. The five classes were mutually exclusive. Within the compatible subset, each conversion event combined its exclusive class with both, so A-to-G and C-to-T event counts could overlap. Other strings could also carry a target-compatible base alongside incompatible changes; their counts were tabulated separately from substitution-compatible numerators. Full-body Q20 all-letter/compatible denominators were 393,868/389,281 for AncBE4max and 53,089/52,623 for NG-ABE8e.

Longer AC-compatible bodies used the same rules against (AC)nGCA, n >8, allowing terminal substitutions; this differs from the eight-dish requirement for unchanged GCA. Zero-support bins were retained. AC7/CCA4 content sensitivities and unit-cost global alignments to AC1–12, CCA1–9 and ACCC1–6 retained tied references and gap ambiguity, without inferring starting material from the nearest match.

For frequencies *p_s_* = *n_s_*/ ∑*_s_ n_s_*, diversity measures were *H* = *−* ∑*_s_ p_s_* log_2_(*p_s_*), *D*_1_ = 2*^H^* and *D*_2_ = 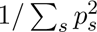. Multivariate hypergeometric sampling matched depth without replacement within identical reference, length, quality and inclusion strata. Depth fractions 0.05, 0.10, 0.25, 0.50, 0.75 and 1.00 of common depth used 200 draws with seed 20261012. At common depth 53,089, the entire NG-ABE8e stratum was used, giving a deterministic point with 107 strings and *D*_1_ = 3.940. AncBE4max had mean richness 72.3 (technical 2.5–97.5% range 65–78) and *D*_1_ = 2.920 (2.886–2.958). These intervals do not quantify between-culture biological uncertainty. Minimum-support sensitivities remained separate from the primary singleton-retaining distributions. Full distributions and singletons were retained. Counts were PCR-weighted; no inferential between-editor test or general performance ranking was made. Quality values were not recalibrated, and neither UMI consensus nor raw-signal rebasecalling was performed. Exact anchors and full-body quality requirements can affect recovery. Content denominators are detailed in Table S1; software versions are listed in “Statistical reporting, visualization and reproducibility”.

#### Capture-barcode recovery and cell aggregation

In both read orientations, the first 220 nt were searched for GCTACGATCCGACTTTCTGCG-N10-CCTTCC-N10CGATG-N10-poly(T), requiring at least eight terminal T bases. The first two N10 segments formed the exact 20-nt C4 bead barcode and the third the 10-nt UMI. No whitelist, tag correction or UMI-neighbour clustering was applied. Bodies were recovered relative to the upstream arm and exact GGGTTAGAGCTAGAAATAGCAAGTTAACCT right boundary and normalized to recorder-forward orientation. Library identities barcode10 and barcode11 were retained separately.

Library 1 bead barcodes were matched exactly in recorded orientation to the vendor bead-to-cell translation, restricted to labels in the delivered expression object, independently of recorder similarity. The restricted translation contained 25,512 beads assigned to 13,743 expression-cell labels, with 7,980 labels linked to multiple beads. Each bead required a unique cell assignment; several beads could map to one label. No mismatch correction, reverse-complement rescue or expression-based reassignment was applied. Unlinked beads remained in the molecular inventory but not the cell cohort. Upstream expression-based cell calling and QC were not inferred from recorder multiplicity.

HQ representatives required body Qeff *≥*20, minimum body-base Q*≥*10 and minimum tag-base Q*≥*20. Within each exact library–bead–UMI family, observed strings entered seed-bounded clusters with Levenshtein budget max(1, floor[0.1 *×* max(length1, length2)]), preserving C/T differences. Representatives were observed readcount-weighted medoids compatible with every member; otherwise the seed was used. Ties used read support, mean body Qeff and lexical order. A representative qualified only if at least one read of that exact string in the family met all quality thresholds. Multiple HQ representatives could remain in one family. No additional read-, family- or recurrence-count cutoff was imposed.

The union of qualifying strings across mapped beads defined each Library 1 cell record set. Library 2 combined strings only within each bead, without expression-based filtering or cross-bead aggregation. No dominant record was imposed, and multiple records did not trigger exclusion. Family-state links were counted separately; UMIs were not deduplicated across beads or equated with independent molecules. Cross-platform matches required identical C4, UMI and complete sequence and could sample the same original molecule. Additional-record recovery was the cell-level union size minus the largest contributing bead set. A sensitivity required at least one supporting family with exactly one retained HQ state; it did not replace primary calls, rebuild the hierarchy or constitute a validated error filter.

The three accepted short-read/ONT matches required identical C4, UMI and complete string, not simply sequence similarity. M1 and M2 each had one ONT read, which met HQ criteria; M3 had two reads, one HQ. M4 failed minimum tag Q20 because its tag minimum was Q18, despite body Qeff 40.551 and minimum body Q36. M4 remained in Source Data but not among the three HQ matches. Such matches may originate from the same molecule on both platforms.

#### Record features, scores and combination hierarchies

Canonical records matched (AC|AT)+GCA. Noncanonical strings and segment boundaries were retained. AT changes, prefixes and contexts of up to two preceding units were extracted from the leading interpretable region, stopping before identified constant-vector sequence. Internal noncanonical fragments supplied exact junction features with up to six flanking nucleotides per side. Eligible edited or noncanonical strings also supplied complete-record features; constant/vector fragments and flagged composites did not provide whole-record lineage evidence.

For a feature present in f of N HQ-positive observation units, its weight was *w* = log_2_[(*N* + 1)/(*f* + 1)] *×{*1 + min[0.75, 0.25 log_2_(1 + *v*)]*}*. Here v was the AT count for change/context/prefix features, noncanonical-fragment length for junction features, or leading-prefix AT count plus noncanonical nucleotide count for complete-record features. Each cell label in Library 1, or bead barcode in Library 2, contributed once to f. Prevalence was calculated for the corresponding cohort, not pooled across libraries.

Record-pair evidence was the largest shared feature weight divided by 1 + *m* + 0.5Δ*u*, where m counted mismatches in the shared interpretable prefix and Δu was the absolute unit-count difference for two canonical records, otherwise zero. For two record sets, B was the maximum record-pair evidence and W the shared eligible exact-record feature weights. Combination evidence was *B* + 0.1 *×* min[*B,* ∑ *W −* max(*W*)], with no additional term for empty W. Missing records incurred no penalty; no one-to-one copy correspondence was imposed. Scores were not calibrated kinship probabilities.

Identical record sets formed one combination node, retaining member observations. Average linkage on 1/(1+evidence) across all combination pairs generated the hierarchy. Combination multiplicity did not weight linkage. Highest-scoring alternatives retained ties within absolute tolerance 10*^−^*^12^; with zero maximum evidence, all alternatives remained unresolved. Candidate summaries were weighted by observation counts. Every observed combination and internal hierarchy edge was retained rather than collapsing subtrees. Child-subtree rotation affected only presentation, not topology or inferred evidence. Internal joins were annotated by records or informative features shared by all descendants, or remained unsupported. These symmetric relations did not assign parents, and layout radius was not time.

Indel-class discrimination compared canonical-only and all-record combinations within the same canonical-positive cohort; indel-only observations were a separate coverage contrast. Exact noncanonical identity was preserved without asserting a verified biological indel event. Examples were selected by sequence structure and support rather than phenotype; the following paragraphs specify the deterministic selection and display criteria.

Figure 4e canonical examples required a shared eight-unit prefix, differences in length and molecular support. Shared-flank examples required one internal noncanonical fragment, at least four units on each side, no more than 18 displayed tokens, at least four variants and an observed canonical comparator. Ranking used flank extent, variant count and support, not phenotype. Every displayed endpoint was present in the expression-linked cohort, and support counted distinct cell labels.

The three-bead aggregation example required four to nine canonical records, at most three observed beads, at least two records added by aggregation and at most 18 AC/AT units per sequence. Eligible cells were ranked by decreasing added-record count, increasing record count and cell label.

The 011 spatial example was the mixed three-unit prefix class with greatest pooled molecular support, selected without coordinates or RNA outcomes. Prefix maps used three, five and eight units. All-zero/all-one classes remained nonspecific background; insufficiently observed prefixes remained unassigned. Simultaneous memberships were preserved rather than forced to a dominant class. The previously explored three-spot crop remained a retrospective illustration, not an independent confirmatory test.

In compact matrices, AC=0 and AT=1 only within interpretable AC/AT regions. Other contiguous fragments occupy an indel-labelled display cell while retaining their exact strings. Comparator dashes are alignment gaps, not called deletions. Trailing blank space lies beyond the observed sequence; no zero-padding extends records artificially. The shared GCA terminal may be omitted visually but is not inferred when absent.

#### Spatial spacer recovery and auxiliary cDNA regions

Spatial tags comprised two 8-nt halves surrounding AGGCCAGAGCATTCGATCCACGTGCTTGAG. The central sequence allowed three edit operations. Concatenated halves were compared with a filtered 16-nt whitelist: uniquely best exact/distance-1 matches formed the primary layer, distance-2 matches were sensitivity data, and tied assignments or split positions were rejected.

The core required body Qeff *≥*20, a recovered UMI and one exact spacer string per pooled spot–UMI key across the two technical libraries. Competing spacer calls remained unresolved; PBS/RTT differences alone did not create spacer conflicts. Deduplication retained 297 states, 15,352 keys and 11,330 spots, with read observations counted separately. No consensus repair, smoothing or imputation was applied, and single-UMI states remained eligible. Spatial combinations used the rarity/complexity score with spot-level prevalence and lower cost for terminal extension than aligned differences. Joins distinguished universal complete-record witnesses, other feature witnesses and unsupported connections. Prefix histories linked to the longest available shorter prefix.

Auxiliary PBS/RTT calls were matched by library/read ID with checks of spot, UMI and orientation; anomalous orientations remained unresolved. Regions required unambiguous full-length alignment and Q*≥*30 at every called base. Repeated-variant support required at least two distinct spot–UMI keys in each compared state. It could rank only existing tied spacer candidates, without changing spacer identities, memberships or spatial/RNA tests. The *≤*1% variant-frequency rule was evaluated as a separate diagnostic. PBS/RTT variation was analysed as auxiliary cDNA evidence. The DNA comparison used separate HEK293T datasets supporting Figure 3, not matched DNA from the spatial specimen.

#### Binary representation and information decomposition

Canonical (AC|AT)+GCA strings were encoded as AC=0 and AT=1, retaining unit count and terminal status. Other exact-GCA strings were aligned by dynamic programming to variable-length A[C/T], with unit mismatch/gap costs and candidate lengths from one to ceil(body_length/2)+2. All globally optimal lengths and traceback transitions were retained, with up to 256 paths per length; none reached this cap. A binary call required agreement across optimal interpretations. Ambiguity, missing observations, deletions, nonbinary substitutions and positions beyond the endpoint remained distinct. Records without GCA retained their literal prefix and unresolved suffix without length imputation.

Structural features required agreement across optimal alignments on operation and allele; equivalent positions remained ranges. Main structural comparisons required at least two pooled spot–UMI keys. Noncanonical model eligibility required unique optimal length, cost *≤*2, at least two interpretable units and no constant-sequence mask. All other records remained in the full ledger and spot sets. Encoding was checked by recon-structing exact observed strings; nonbinary substitutions were not relabelled as indels. Compact display gaps were not called deletions, and no sequence was zero-padded beyond its observed end.

For the 102 canonical spatial records, information was decomposed as *H*(*F*) = *H*(*P*) + [H(P,L)-H(P)] + [*H*(*F*) *− H*(*P, L*)], where F is the full record, P the visible eight-unit prefix and L full length [46]. Equal endpoint weights and 14,924 pooled canonical spot–UMI weights were analysed separately. The 25 records shorter than eight units retained visible boundaries; truncated records had censored terminal length. Prefix-only keys excluded full length, state identity and downstream alleles. Length increments used *H*(*P, L*) *− H*(*P*), not H(L); conditional pattern entropy was *H*(pattern, length) *− H*(length).

Structural information compared the same 73 supported records with exact identity hidden or available, preserving homology masks and length. Raw-identity upper bounds on all model-eligible endpoints were separate. C4-level contrasts used unordered canonical-record sets within the same beads without pooling libraries or inferring cell identities.

#### Independent-character and ordered-extension models

Both models used the same 183 eligible records, scores and candidate pool of ordered pairs of distinct endpoints. At least two jointly known binary units were required; missing positions supplied neither agreement nor editing evidence. Both primary models prohibited observed 1*→*0 reversals. Common cost counted overlapping 0*→*1 changes, three times overlapping 1*→*0 changes, new 1s beyond a known source endpoint, and 0.5 times the minimum separation of length intervals. Each symmetric-difference structural feature added one only when observable in both records. Unknown structural state was not absence; costs were not transition probabilities.

The independent model allowed changes at all observed positions and either direction of length change. The ordered model additionally required interval noncontraction and prohibited new 0*→*1 changes before max(0*, L*_source_ *− W*), using exploratory W=8. Initial-register substitutions had no imposed left-to-right order; equal-length transitions and unsampled intermediates were allowed. Only admissibility masks differed. Incoming candidates within one cost unit of the best score, including ties, were retained. Endpoints without admissible parents remained unplaced; strongly connected components retained cycles, and only their quotient graph was acyclic. No forced root-adjacent fallback or unique parent was inserted.

Four full-record configurations crossed model type with structural identity on/off on 183 endpoints. Two eightunit-prefix configurations and their full-record references used 102 canonical endpoints. Censored prefixes had length interval [8,*∞*), without downstream terminal, original length, raw ID or hidden alleles. Independent scalar calculations checked all scores and candidate masks; synchronized position-column and metadata relabelling left the independent-model result unchanged.

#### Perturbations and simulated genealogies

Sensitivities used windows of seven/nine units, one allowed archived-site change, contraction-permitted ordering and a shared one-reversal allowance. Random masking of 5%, 10% or 20% of known binary positions preserved order. Loss of 10%, 20% or 40% of pooled spot–UMI keys used 50 repeats per setting, retaining source state calls and structural definitions. A record survived with at least one sampled key. Candidate Jaccard and baseline retention were conditional on survival; coverage was reported separately. Technical libraries were also evaluated separately.

Simulated genealogies were balanced binary trees of depth five with 32 terminal nodes and an unconverted eight-unit root. A branch extension added one unconverted unit, or two with conditional probability 0.2. Cto-T edits were sampled before extension at the scenario probability in the active window, or at the specified archived-site probability, and again after extension at half the scenario probability in the new active window. Initial and existing sites did not receive deterministic branch codes. Unless specified otherwise, internal-node sampling probability was 0.55.

| Scenario | Explicit scenario settings |
| --- | --- |
| Ideal terminal editing | Window 8; edit probability 0.18; extension probability 0.65 |
| Wider active window | Window 12; edit probability 0.07; extension probability 0.85 |
| Saturation/homoplasy | Edit probability 0.75; extension probability 0.5; positional pre-extension multipliers 0.35–1; internal-node sampling probability 0.2 |
| Contraction/re-editing | Delete one uniformly chosen unit after editing with probability 0.18; archived-site edit probability 0.06 |
| Scenario | Explicit scenario settings |
| Error/dropout | Mask 10% of positions; bit-flip probability 0.025; lose 20% of otherwise observed non-root nodes; censor the last two units with probability 0.10 when length exceeds four |

For every scenario except the wider-window case, the window was eight units. Default edit and extension probabilities were 0.18 and 0.65; the wider-window and saturation values in the table replace these defaults. Archived-site editing, contraction, masking, bit flips and observation loss were zero unless specified. Internal-node sampling was 0.55 except for saturation (0.2). Contraction was evaluated after post-extension editing only for lengths greater than three. Saturation multiplied pre-extension editing at zero-based position k by 0.35 + 0.65 *×* ((k mod 4)/3). Observation errors were applied before position masking; terminal censoring followed both. These probabilities define the simulation scenarios. Complete per-scenario values are also provided in the Simulation_parameters resource. Twenty genealogies per scenario used seeds 2026092713+1000*×*scenario_index+replicate_index. A separate timing pilot used pilot_seed+9,000,000+scenario_index, with no pilot outcome used for tuning. Latent nodes retained parent identities and exact true sequences; all 100 genealogies and observed datasets were saved and hashed before inference. Both models received the same noisy observations, and no experimental inferred tree supplied ground truth.

Both models received identical noisy observations. True relationships were defined from saved ancestry and, separately, the nearest sampled ancestor. Metrics were true-relation retention, false-pair admissibility, near-candidate precision, unique-best wrong-relation fraction and coverage; they evaluated candidate relations rather than full-tree accuracy or recorder architectures. Roots remained in target-coverage and unique-best denominators; ancestor-retention denominators included only targets with actual sampled ancestors. Experimental inferred trees were not used as truth. All replicate outcomes, including failures and non-gains, were retained; separate timing pilots did not tune the test simulations.

Masking and pooled-key-loss sensitivities used seed 2026092711. The timing pilot seed was 2026092712.

#### Spatial and RNA association tests

Spatial tests used the 11,330 core-bearing spots and 18,633 undirected first-ring edges joining measured centres within 75 full-resolution pixels. Missing spots were not replaced by distant neighbours. Groups required at least three positive and three negative spots, yielding 114 tests. Whole membership vectors were jointly permuted 9,999 times (seed 20260924) within 4*×*4 tissue blocks, RNA-depth quartiles and log2 core-state-count bins: 147 strata, including 13 singletons. This preserved group sizes and between-group co-occurrence within strata.

Spatial P values were one-sided, (1+null counts at least observed)/(9,999+1) [47]. A planned 99,999-draw refinement for pilot P*≤*0.005 was not triggered. Benjamini–Hochberg correction covered all 114 tests [48]. RNA tests required at least ten positive and ten negative spots, giving 57 groups and nine programmes; all 513 tests used two-sided deviations from the exact stratum-weighted expected score sum under the same permutations and were corrected together. Permutation ranges describe conditional null distributions, not biological confidence intervals.

Programme scores used the delivered log1p expression matrix after scaling each spot to the median library total of 744 counts. Available marker expression was standardized across all 28,739 tissue spots using sample standard deviation, clipped to [-5,5], and averaged within programmes. Symbols were matched case-insensitively; Mx1 was unavailable, leaving six interferon markers. Programmes covered epithelial, EMT/invasion, hypoxia/glycolysis, proliferation, interferon, immune, myeloid, fibroblast/ECM and vascular expression; exact gene lists accompany Source Data.

Scores were regressed by ordinary least squares on an intercept, log1p total counts, log1p detected genes, standardized x/y coordinates, x^2^, y^2^ and xy. Residuals were centred and scaled to unit sample standard deviation across all tissue spots, then analysed in the fixed core-bearing cohort. Cluster dummy variables were not included. Scores represent spot-level marker programmes, not cell-type proportions. Molecular groups were fixed before these tests, but the tissue had been explored previously; analyses were retrospective.

#### Paired spatial model comparison

Encoded records, candidate relations, group memberships and examples were fixed before loading new spatial-test outcomes. Prefix maps used three, five and eight units; insufficiently observed records remained unassigned. All-zero/all-one classes remained nonspecific background. Every class and simultaneous spot membership were retained. The 011 example was selected by greatest pooled support among mixed three-unit prefixes, without coordinates or RNA outcomes. Prefix nesting was descriptive rather than validated clonal ancestry.

A symmetric model relation required at least one near-best directed relation and a shared AT or usable structural feature; informative exact identity could also support cross-spot association. Pure design matches and pairs of fully saturated canonical strings were excluded as specific witnesses in both models. All 297 records remained in spot bundles. Both models used the same 11,330-spot universe and 18,165 eligible first-ring edges, with at least one eligible record at each endpoint and the same 75-pixel distance threshold.

Three aggregations were any supported cross-spot pair, mean support across all pairs, and mean support weighted within spots by log(1+nUMI). Unmodelled records supplied no positive support but remained in denominators. Each of 9,999 permutations shuffled whole record/UMI bundles within spatial block, RNA-depth bin, exact total record count, exact eligible-record count and UMI category capped at four. Each assignment was shared by both models and all aggregations.

Model-specific enrichment used one-sided (1+exceedances)/(1+permutations) tests [47]. Direct orderedminus-independent differences were centred on the paired-null mean and tested two-sided. Benjamini– Hochberg correction jointly covered nine tests: two models and their difference under three aggregations [48]. Central 95% ranges were null ranges, not confidence intervals. Both models used the same sequence-defined groups.

Paired spatial permutations used seed 2026092714.

#### Statistical reporting, visualization and reproducibility

Counts retained assay-specific units and inclusion sets. Record-combination and support summaries were descriptive, without inferential tests. The dish-level two-test Holm family, 114 spatial tests, 513 RNA tests and nine paired-model tests remained distinct. Exploratory sampling diagnostics were identified separately. Fixed seeds and complete outputs were retained, including unresolved candidates and negative results.

Spatial maps retained simultaneous record memberships at measured coordinates registered to the tissue image.

Versioned manifests link input files, scripts, configurations, Source Data and figures. Software environments and computational checks are described in this subsection. Analysis tables retain exact sequences, raw orientations, paired or one-sided recovery tiers and molecular keys. Qeff summarizes base-error probabilities, rather than the correctness probability of an entire sequence.

Measured *x*_hires_ and *y*_hires_ were full-resolution coordinates multiplied by 0.18424689083371718 and registered to a 2000*×*2000 tissue image. Matplotlib RegularPolygon hexagons [49] used circumradius 4.5140488 display pixels, equal to 0.49 times the median nearest-neighbour distance of 9.2123445 pixels across all 28,739 tissue spots. The hexagons are display glyphs on a Cartesian array. Whole-section plots used equal aspect, limits -0.5 to 1999.5 and downward y matching the image. Coordinates were not reflected, moved, smoothed or interpolated. H&E overlays used uniform alpha 0.40; scale rulers were image pixels without micrometre calibration.

Pale grey represented measured spots without core evidence; a second grey distinguished other core-bearing spots. Mixed membership had its own category where needed. Presence, molecular-count and mixed-membership panels retained separate units and scales. Analysis adjacency was defined in full-resolution coordinates at 75 pixels, not by the display-hexagon radius.

Figure 6 spatial effects were log_2_(observed/null-mean edges + 0.125), plotted against *−* log_10_(*P*), with a nominal P=0.05 line. In Figure S10, the no-enrichment reference was log_2_(1.125), not zero; groups with both null mean and observed count zero were N/A. Variable-prefix, exact-indel, shared-indel and exact-regular groups used teal, vermilion, muted purple and blue. R1/R2/R3 denoted ACAT/AT/ACACAT entry prefixes.

RNA display values were group-mean programme scores minus exact stratified-null expectations, with limits ±0.3 for Figure 6 and ±0.8 for the complete Figure S10 matrix. The eight-state example used gold for its dominant exact state, navy for other repeatedly supported states and open glyphs for singletons; its sequence view used pale teal for AC, navy for AT and grey for GCA. Colour and crop choices did not change tested membership or multiple-testing families.

Library 1 processing used consensus_core.py, build_evidence_tables.py and analyse.py. Family representatives were selected from observed strings using the eligibility and tie-breaking rules above. Vectorized record and combination scores were checked against direct calculations. Library 1 feature prevalence was recalculated for expression-cell labels; Library 2 retained its bead-barcode weights and hierarchy. All nodes, edges, member observations and tied highest-scoring candidates were preserved.

Quantitative encoding was checked for exact-string reconstruction, including all optimal-length interpretations and terminal-unresolved suffixes. A separate scalar implementation verified scores and admissibility masks in all eight model configurations. Synchronized column/metadata relabelling left the independent model unchanged. Input, configuration, encoding, simulation truth and paired-null outputs were versioned separately from upstream expression processing.

Basecalling records specified Dorado v7.8.3 and sequencing reports MinKNOW v25.03.7, with demultiplexing within MinKNOW. Main-figure reads used SUP v4.3.0; the two editor FASTQ inputs recorded dna_r10.4.1_e8.2_400bps_hac@v4.2.0. The Library 1 environment recorded Python 3.12.14, NumPy 2.5.2 and pandas 3.0.1. Its SciPy version was not recorded; the separate binary-model environment recorded SciPy 1.18.1. Runtime versions and dependency requirements are listed separately.

## Supplemental information

Self-extending DNA recording with stochastic and programmed writing

Figures S1–S10 and their legends accompany the main manuscript. Table S1 and machine-readable source tables are provided separately; detailed methods are integrated into STAR Methods.

---

**Figure S1.**
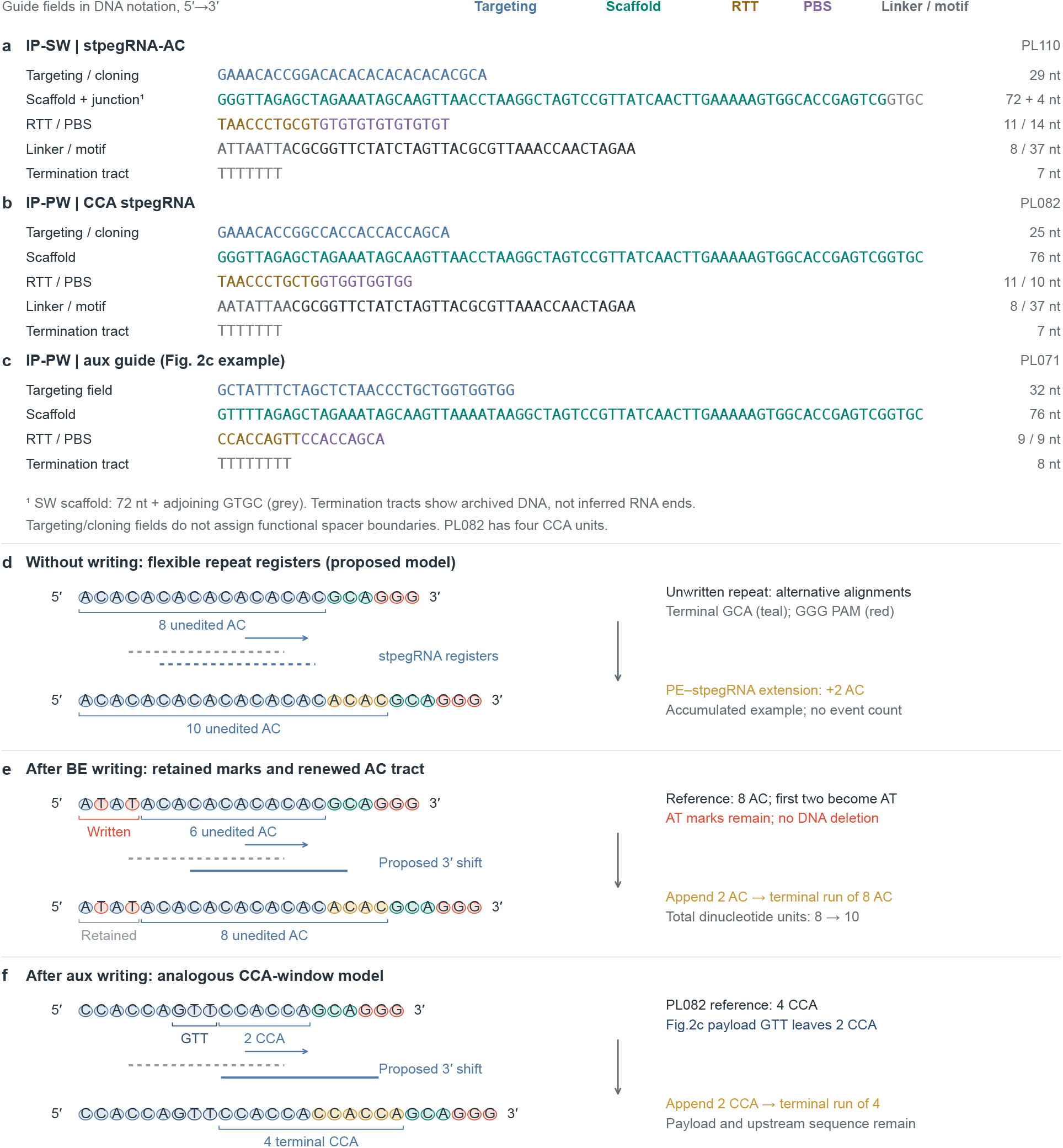
Guide sequences and repeat-window renewal, related to Figures 1 and 2. a, IP-SW stpegRNA-AC sequence fields, with the complete local DNA window retained from the targeting/cloning field through the termination tract. The 72-nt scaffold field is coloured teal; the adjoining GTGC is shown separately in grey. b, The four-CCA IP-PW stpegRNA construct. The 21-nt 3*^′^*extension comprises an 11-nt RTT and a 10-nt PBS, according to the construct annotation. c, The aux guide used for the sequence model in Figure 2c, with its 9-nt RTT and 9-nt PBS; this AAC-template example corresponds to GTT in the recorder-forward DNA. Bases are displayed in DNA notation and read 5*^′^* to 3*^′^*. Blue denotes targeting/cloning fields, teal scaffold, ochre RTT, purple PBS, slate linker and dark grey the stabilizing motif. Other construct context and termination tracts are grey. The separate field labels retain each guide’s actual sequence; targeting/cloning field lengths do not establish functional spacer lengths. Complete sequences for all nine aux constructs are provided in the construct resources. d, An unedited AC repeat tract can accommodate alternative repeat registers in the proposed PE–stpegRNA model. The lower row illustrates an accumulated two-AC extension; the arrow summarizes the accumulated extension. e, Eight-unit design example. Conversion of the first two AC units to AT leaves a terminal run of six unedited AC units without deleting DNA. The proposed subsequent addition of two AC units restores an eight-AC terminal run while retaining the AT prefix, giving ten total dinucleotide units. f, An analogous model for the separate CCA-based IP-PW design. The GTT example from Figure 2c places two CCA units downstream of the payload. Appending two CCA units would restore a four-CCA terminal run, with the payload and upstream sequence retained. The four-CCA reference is the PL082 construct. Ochre bases mark illustrative additions, vermilion T bases mark the BE example, navy denotes the payload, teal terminal GCA and vermilion GGG the stpegRNA PAM context. DNA is shown in recorder-forward orientation, 5*^′^* to 3*^′^*. Register bars denote repeat positions. The proposed routes and restoration targets remain mechanistic models requiring experimental validation.

**Figure S2.**
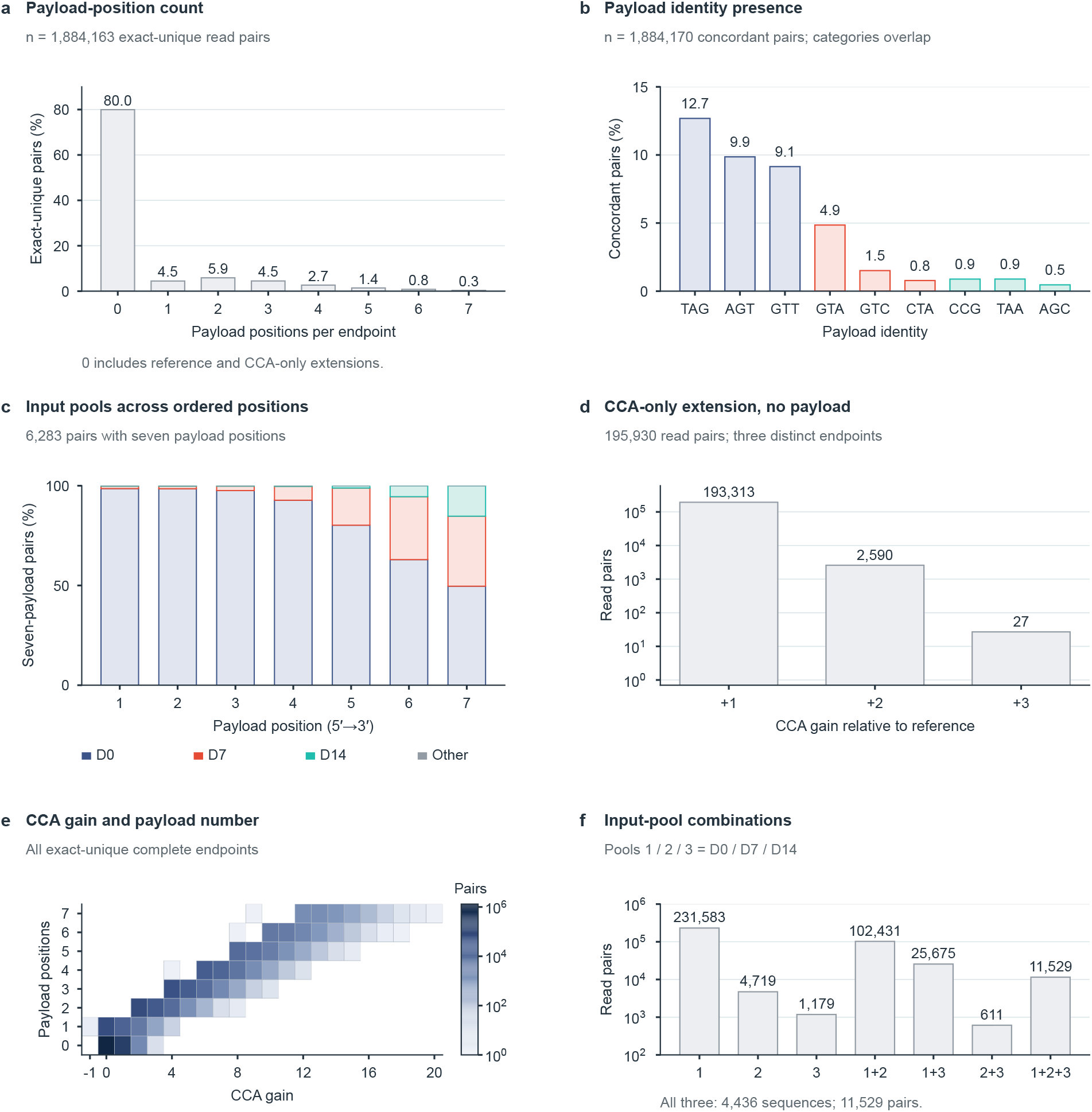
Complete programmed-writing endpoint distributions, related to Figure 2. a, Parsed payload-position counts among 1,884,163 exact-unique complete-body read pairs. Counts for depths 0-7 are 1,506,430, 84,042, 110,889, 84,758, 50,167, 26,524, 15,070, 6,283. The zero bin contains the reference and all CCA-only extensions. b, Presence of each specified triplet among 1,884,170 mate-concordant complete-body pairs. A pair contributes once to each identity it contains; bars overlap in membership. Colours match Figure 2d. c, Input-pool composition at each observed 5*^′^→*3*^′^* payload position among 6,283 read pairs with seven payload positions (5,844 distinct exact-unique bodies). Each position uses the same denominator. Stacks run from bottom to top in D0, D7, D14 and other order. Colours indicate sequence-identity membership in an input pool, not an inferred incorporation time. The grey other category retains one pair with an outside-pool triplet at position 7. d, All payload-free CCA expansions: +1 CCA, 193,313 pairs; +2 CCA, 2,590 pairs; +3 CCA, 27 pairs. The three bodies sum to 195,930 pairs. The count axis is logarithmic. e, Joint count distribution of CCA-register gain and payload-position count among all exact-unique complete endpoints. CCA gain is measured relative to the four-CCA reference after separating parsed payloads. Coloured cells show read-pair counts on a logarithmic scale, and white cells contain zero pairs. f, Exclusive membership combinations of the three specified input pools (1=D0, 2=D7, 3=D14): 1: 231,583; 2: 4,719; 3: 1,179; 1+2: 102,431; 1+3: 25,675; 2+3: 611; 1+2+3: 11,529. Six additional exact-unique payload-bearing pairs contain triplets outside these nine specified identities and are retained in the source table, outside the seven displayed combinations. The all-three-pool category comprises 4,436 distinct sequences and 11,529 read pairs. The count axis is logarithmic. Data and statistics. All panels use the existing complete-endpoint catalogue from the CCA-mixed sample harvested at D21 after payload transfections on D0, D7 and D14. There are 2,220,942 synchronized input pairs, 1,884,170 mate-concordant complete-body pairs and 42,648 distinct complete bodies. Payload statistics use the existing exact-unique parsing rule and the nine specified identities; CCA is assigned to the repeat register. Counts are descriptive PCR-weighted read-pair occurrences. Source tables link to the complete catalogue of counted endpoints. The IP-PW endpoint data were obtained from one sample of HEK293T cells (biological n = 1).

**Figure S3.**
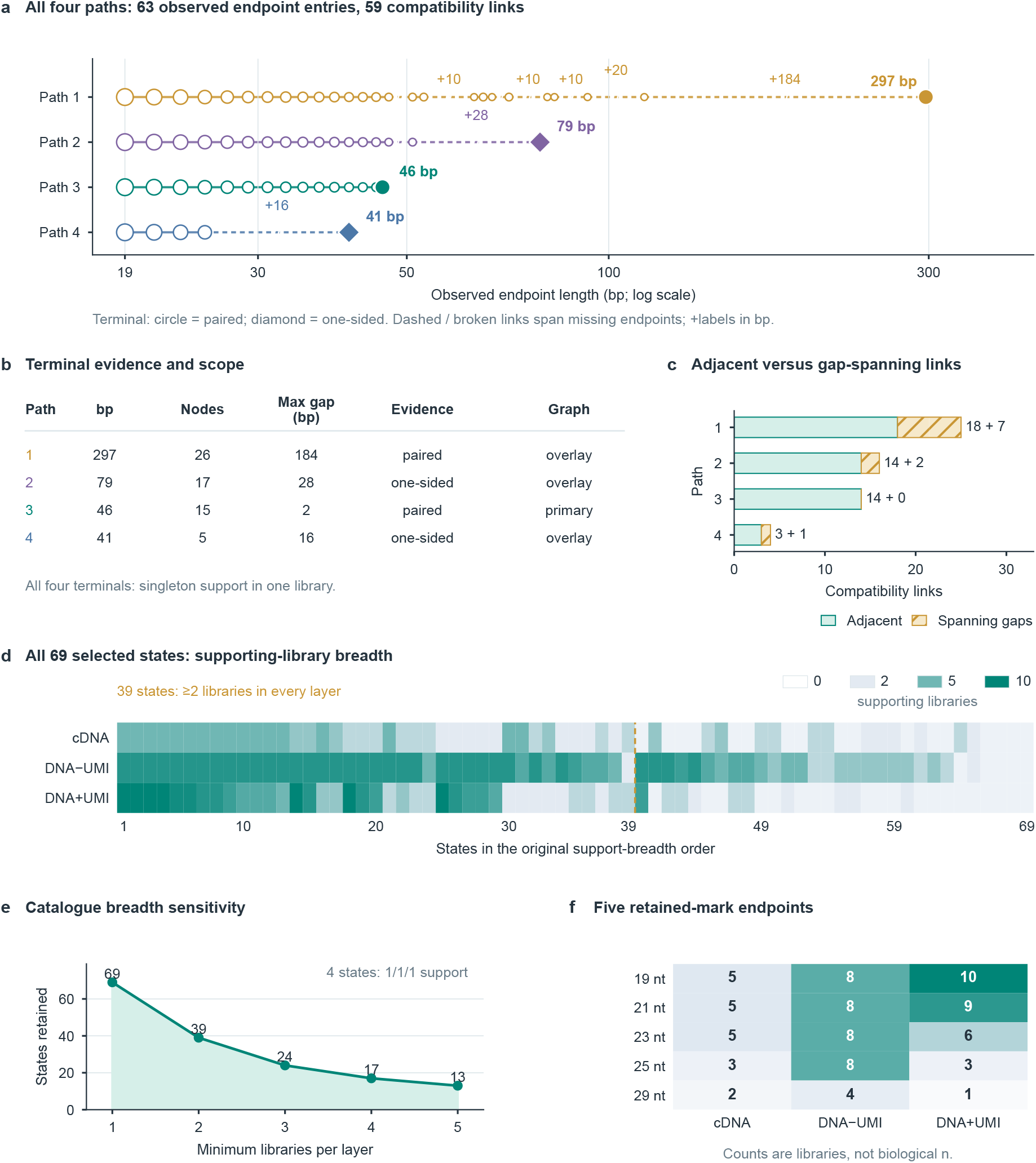
Compatible endpoints and cross-library support, related to Figure 3. a, Four retained-sequence-compatible paths contain 63 observed endpoint entries and 59 compatibility links; some endpoints are shared between paths. Node area scales with log-transformed UMI support. Terminal circles denote paired full-body evidence and diamonds one-sided evidence. Dashed/broken links span missing intermediate endpoints; numeric annotations report gaps of at least 10 bp. Links denote sequence compatibility rather than tracked transitions, time, editing cycles or ancestry. b, Terminal length, observed-node count, maximum length gap, terminal evidence type and primary-versus-overlay membership for each path. The 297-, 79-, 46and 41-bp terminals each have singleton support in one library; only the 46-bp terminal belongs to the primary graph. c, All 59 links classified as joining adjacent observed endpoints or spanning missing endpoints. Printed counts report both categories, including zero gap-spanning links in path 3; maximum gaps are given in b. d, All 69 selected expanded states across transcriptome-derived cDNA, non-UMI targeted DNA and UMI-tagged targeted DNA, ranked by minimum library breadth, then mean breadth and sequence length. Cell colour is log2(1 + supporting libraries). All three layers were required for catalogue admission. The dashed boundary marks 39 states with at least two libraries in every layer. e, Catalogue-retention sensitivity to requiring at least one to five supporting libraries simultaneously in each layer: 69, 39, 24, 17 and 13 states. Four states have exactly 1/1/1 library support. f, Supporting-library counts for five prespecified 19-, 21-, 23-, 25and 29-nt retained-mark endpoints. Matrix entries count supporting libraries. Across-layer support describes library breadth within the selected catalogue, not a biological-replicate statistic. Exact sequences and counts are provided in Source Data. Non-UMI DNA libraries 01–04 correspond to BE + stpegRNA and libraries 05–08 to PE + BE + stpegRNA, with four independent dishes per condition.

**Figure S4.**
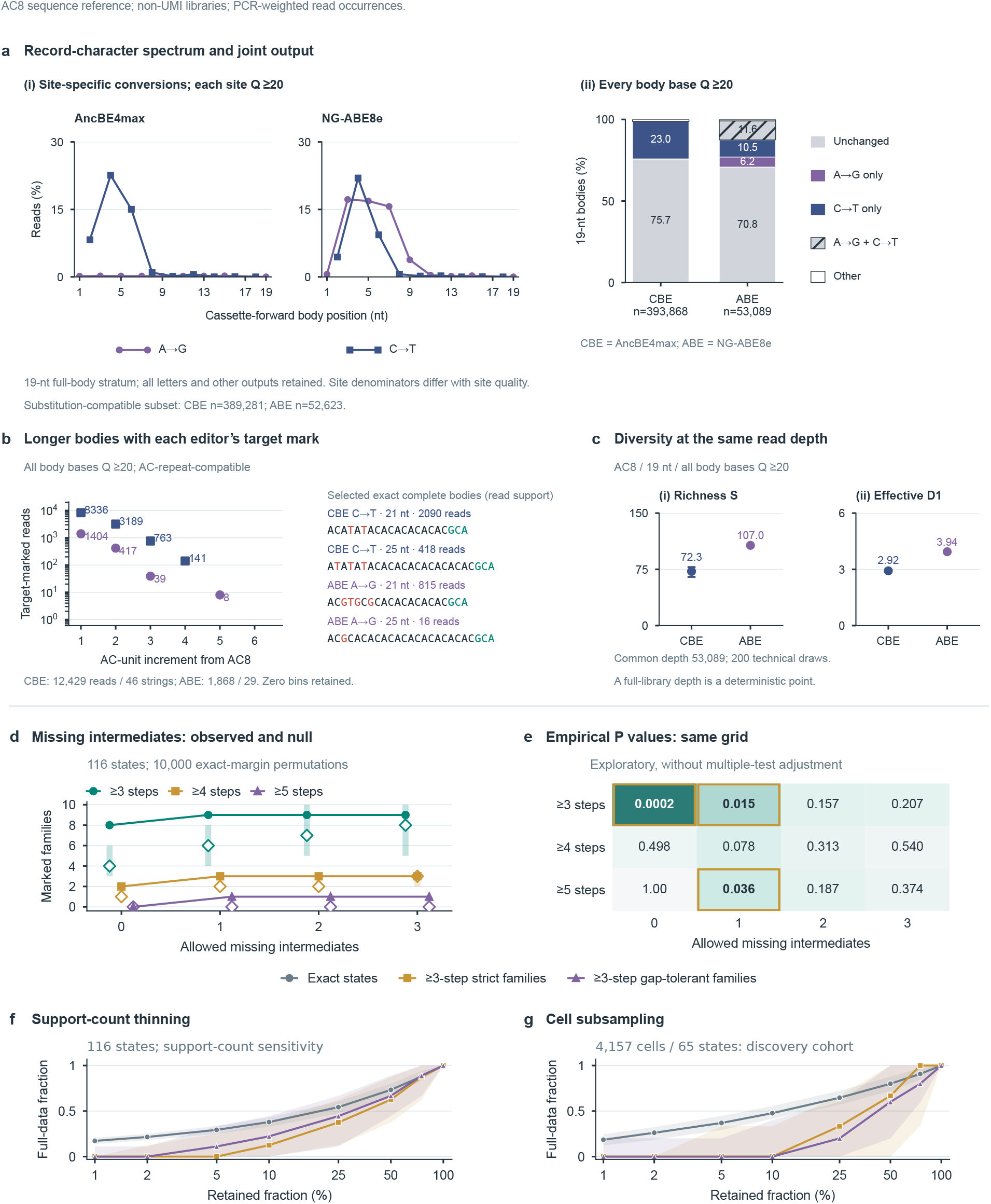
Editor character spectra and observation completeness, related to Figure 3. a, Position-specific A-to-Gand C-to-T-compatible fractions among complete AC8-length bodies, in cassette-forward coordinates. Each site denominator includes all alleles meeting site Q*≥*20; other body positions need not meet Q20 for this layer. The accompanying joint-state composition requires Q*≥*20 at every body base and retains the five mutually exclusive classes unchanged, A-to-G only, C-to-T only, both and Other; n=393,868 AncBE4max-condition and 53,089 NG-ABE8e-condition read occurrences. The Results use the substitution-compatible subset (n=389,281 and 52,623), comprising the first four classes. Within that subset, at least one A-to-G change equals A-to-G only plus both; at least one C-to-T change equals C-to-T only plus both. NG-ABE8e A-to-G is 9,478/52,623 (18.01%) in the subset and 9,478/53,089 (17.85%) when those same compatible-event members are normalized to the all-letter display; AncBE4max C-to-T is 90,750/389,281 (23.31%) and 90,750/393,868 (23.04%), respectively. Other outputs remain in the all-letter denominator and may also contain a target-compatible base alongside incompatible changes; their overlapping conversion counts are retained separately in Source Data. b, Target-marked AC-repeat-compatible bodies with Q*≥*20 at every body base, longer than the (AC)8GCA sequence reference (19 nt) and grouped by repeat increment. AncBE4max-condition C-to-T-marked outputs total 12,429 occurrences and 46 distinct complete bodies; NG-ABE8e-condition A-to-G-marked outputs total 1,868 occurrences and 29 bodies. Sequence examples show the most-supported exact target-marked body at 21 and 25 nt in each condition, with lexicographic tie-breaking; an absent example is displayed as no observed support. Coloured bases indicate changes relative to the complete (AC)nGCA reference, with the terminal segment shown separately. Compatibility allows reference A-to-G and C-to-T substitutions, including terminal positions, with no insertions, deletions or other substitutions. Counts quantify PCR-weighted read support. c, All-letter AC8-length (19-nt) outputs with Q*≥*20 at every body base, compared at matched read depth for observed sequence richness S and effective Shannon states *D*_1_ = 2*^H^*. The common depth is 53,089 occurrences. AncBE4max points denote the mean and bars the technical 2.5-97.5% range of 200 without-replacement draws (richness, 72.3, range 65-78; effective states, 2.920, range 2.886-2.958). This depth equals the entire NG-ABE8e layer, whose technical distribution is deterministic; it is shown as a point without an error bar (107 strings; effective states, 3.940). These panels describe one non-UMI sequencing library per editor configuration, processed from the complete merged FASTQ input. The editors were selected after comparison of published studies; no experimental sample or sequencing library was excluded. Read occurrences are PCR weighted; technical subsampling ranges are separate from biological uncertainty. d, Missing-intermediate sensitivity in the 116-state cross-library catalogue detected across all three library types. Filled symbols show observed marked families spanning at least three, four or five nominal +AC steps; open diamonds and vertical bars show the median and central 95% range from 10,000 exact-margin permutations. Allowing one missing intermediate admits the 19*→*21*→*23*→*25*→*29-nt family, with the 27-nt intermediate unobserved, and increases the maximum marked nominal depth from four to five. e, One-sided upper-tail empirical P values for the same twelve settings. P<0.05 cells are outlined; the exploratory grid has no multiple-comparison correction. Full-precision P values are in Source Data. f, Cross-layer support-count thinning. g, Uniform cell subsampling of the 4,157-cell / 65-state discovery call set. Lines show medians and shading central 95% ranges from 1,000 resampling repeats at each subsampled fraction; the full-data endpoint is deterministic. Exact-state fractions are normalized to their full-data counts; strict and gap-tolerant *≥*3-step family counts are normalized by 8 and 9 for f, and by 3 and 5 for g. In the cell-subsampling discovery set, 4,166 of 6,137 cell–state links (67.9%) have one observed UMI; 2,482 of 4,157 cells (59.7%) contain only single-UMI states. At 10% cell sampling, the median retained-state fraction is 47.7% and the median number of *≥*3-step marked families is zero. The discovery cohort is separate from the final 12,302-cell aggregation cohort. Thinning measures recovery within the observed catalogue; compatible histories describe sequence relationships. The editor libraries, the 116-state diagnostic and the cell-subsampling set use separate observational units and denominators. Editor character spectra in a-c describe AC8-length or AC-repeat-compatible subsets; their full admission and content-class ledgers, Q sensitivities and exact frequencies are provided as Source Data. AncBE4max was selected for the binary implementation because its C-to-T-directed output aligned with the AC/AT alphabet.

**Figure S5.**
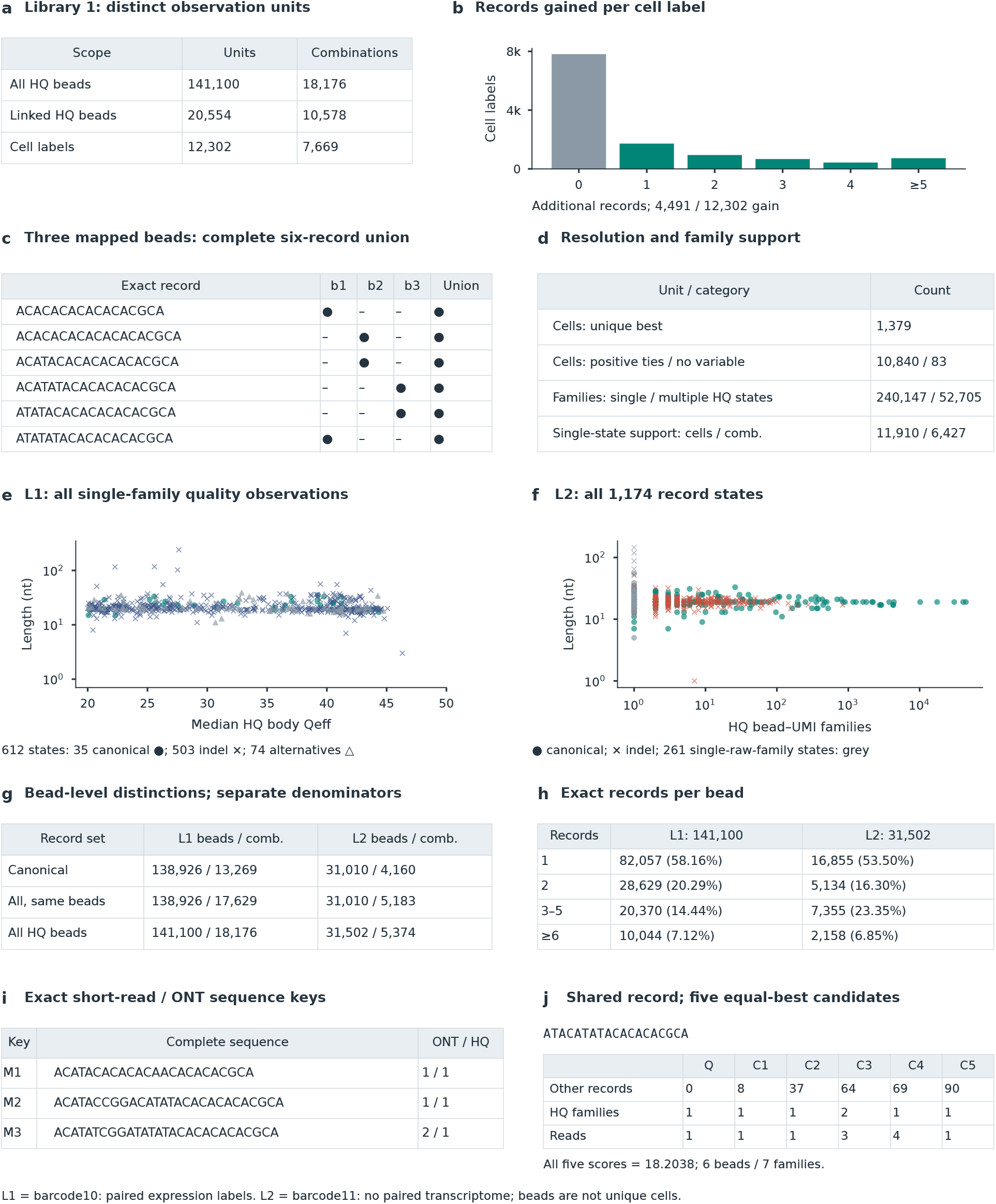
Capture-library units and molecular support, related to Figure 4. a, Cohort restriction and the observation unit are shown separately: the full molecular inventory contains 141,100 HQ bead tags and 18,176 combinations; 20,554 expression-linked HQ beads define 10,578 combinations; aggregation into 12,302 expression-cell labels gives 7,669 combinations. The cell cohort comprises expression-linked beads after aggregation. b, Distribution of additional exact records after bead-to-cell aggregation, relative to the largest record set recovered from any single bead mapped to that cell. Bins are 0, 1, 2, 3, 4 and at least 5 additional records. Overall, 4,491 of 12,302 labels gain records. c, An observed three-bead example (CELL5267_N3). Each mapped bead contributes two distinct records, and their union contains all six. Complete record sequences and the six-by-four bead/union membership matrix are shown. Filled circles denote observed membership; dashes denote absence from the respective observed set. Exact bead keys and state IDs are retained in Source Data. d, Cell-weighted candidate relation classes: 1,379 unique-best labels, 10,840 positive-evidence ties and 83 labels with no variable shared evidence. The latter two sum to 10,923 labels without a unique best candidate. The retained evidence contains 292,852 distinct bead–UMI families: 240,147 with one HQ state and 52,705 with multiple HQ states. Family-support sensitivity used these molecular observations. Retained calls give 12,302 cell labels and 7,669 combinations; restricting support to families with a single HQ state gives 11,910 labels and 6,427 combinations. Library 1 is barcode10 and is the only capture library linked to the paired expression matrix. e, All 612 Library 1 exact states observed in one raw bead–UMI family and supported by at least one HQ read. There are 538 selected representatives (35 canonical circles and 503 indel-class crosses) and 74 within-family alternatives (triangles); 601 states have one raw read. Length is logarithmic; the horizontal axis is median HQ body Qeff. The all-tag inventory includes states outside the expression-cell cohort. The existing HQ criteria are body Qeff *≥*20, minimum body Q *≥*10 and minimum tag Q *≥*20. Base-call quality is not a whole-sequence correctness probability. f, All 1,174 retained Library 2 exact records by distinct supporting HQ representative bead–UMI families and nucleotide length, both logarithmic. Circles denote canonical AC/AT records and crosses indel-class records. The 261 representatives found in one raw family are grey; all other points retain class colour. Exact noncanonical strings remain in Source Data. g, Canonical-only, all-record/same-bead and all-record/all-HQ-bead combination counts in each library. Within the canonical-positive cohort, all records add 4,360 distinctions in Library 1 and 1,023 in Library 2. Including indel-only bead groups adds a further 547 and 191 combinations, respectively. These comparisons distinguish within-cohort discrimination from additional coverage; they are bead-level counts, including Library 1 beads outside the linked expression cohort. h, Full distribution of retained exact records per HQ bead barcode in both libraries. Counts and percentages use denominators 141,100 and 31,502. Categories are one, two, three to five and at least six records. These multiplicities describe recovered record sets per bead; integrated copy number and doublet status were not measured. i, Three exact cross-platform molecular matches M1–M3. Each requires identical C4 bead tag, UMI and full nucleotide sequence in short-read and ONT evidence. Each identical sequence key is printed once, including noncanonical fragments and terminal GCA; the ONT and HQ counts remain separate. M1/M2 each have one ONT read, which is HQ; M3 has two ONT reads, one HQ. The additional M4 key fails the existing minimum-tag Q20 criterion (Q18), despite body Qeff 40.551 and minimum body Q36; its sequence has 69 HQ representative families across 42 other C4 groups. Full M1–M4 keys and quality values remain in Source Data. Exact-key matches may concern the same original molecule and are not independent replication. j, Six Library 2 bead groups share exact S record ATACATATACACACACGCA, supported by seven HQ representative families. The query contains S alone; the five candidates contain 8, 37, 64, 69 or 90 additional records and all tie at the highest score, 18.2038. Family/read rows refer to S in each group, whereas additional-record counts describe the remainder of each observed combination. Q and C1–C5 map to complete bead keys in Source Data. Missing records remain unknown, leaving all five candidate relations tied. Library 1 (barcode10) alone has paired expression-cell labels. Library 2 (barcode11) is an independent biological replicate under the same experimental conditions, with barcode-targeted cDNA sequencing and no paired transcriptome; its observation unit is the capture bead barcode. The complete Library 1 hierarchy remains in Figure 4b. Library 2 hierarchy nodes, display order and observation membership are provided in Source Data.

**Figure S6.**
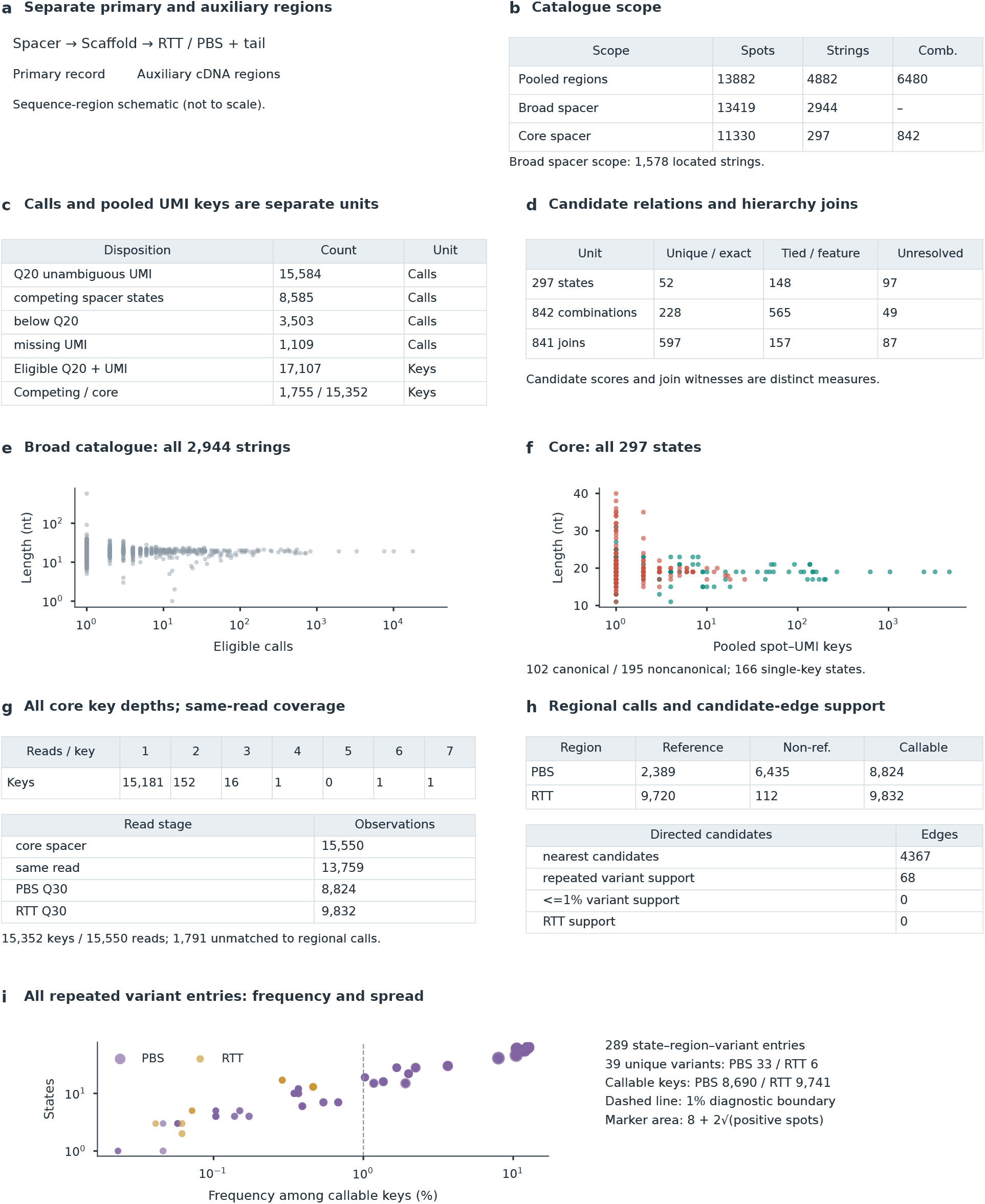
Spatial record scope and auxiliary evidence, related to Figure 5. a, The scaffold-upstream spacer defines the primary record; downstream RTT/PBS sequence is catalogued separately as auxiliary cDNA evidence. The region schematic is not to scale. b, Counts under pooled-region, broad-spacer and core-spacer scopes. “Strings” denotes exact sequence strings, including the 297 core states; “Comb.” denotes record combinations. The broad catalogue includes 1,578 located strings. The dash indicates a combination count not reported here, not zero. Each scope uses its own inclusion criteria and denominator. c, Disposition of 28,781 located calls before duplicate removal: 15,584 Q20 calls at unambiguous recovered UMI keys, 8,585 competing-state calls, 3,503 below-Q20 calls and 1,109 calls without a recovered UMI. These call counts differ from pooled-key and core read-observation counts. The same pooled spot-UMI unit throughout: 17,107 eligible keys, 1,755 competing-state keys left unresolved and 15,352 core keys. Calls and pooled keys are retained as separate counting units. d, Candidate status for 297 states (52 unique best, 148 tied best and 97 without shared evidence), for 842 combinations (228, 565 and 49, respectively), and witness classes for 841 tree joins (597 complete-record, 157 feature and 87 unresolved). These categories summarize candidate scores and the evidence supporting tree joins. e, All 2,944 broad spacer strings, comprising 59,031 eligible calls, plotted by call support and length; both axes are logarithmic. f, All 297 core states by length and pooled spot–UMI-key support. Canonical states match the retained AC/AT-repeat-plus-GCA grammar (102 states); the other 195 states are labelled noncanonical. A total of 166 states have one pooled key. Noncanonical status alone does not establish an insertion or deletion. g, Read-depth distribution for all 15,352 core pooled keys, representing 15,550 read observations; 15,181 keys have one read. The exact counts, including zero at depth five, are tabulated. Physical same-read matching and regional callable observations: 15,550 core spacer read observations, 13,759 matched regional observations, 8,824 PBS Q30 observations and 9,832 RTT Q30 observations. The PBS and RTT callable sets may overlap. There are 1,791 unmatched core observations; three matched observations lacking the expected opposite orientation remain unresolved. Exact depth counts and regional read counts are tabulated. h, Reference and non-reference observations within each region-specific callable-read denominator: PBS, 2,389 and 6,435 of 8,824; RTT, 9,720 and 112 of 9,832. Of 4,367 existing directed nearest candidates, including unique-best and tied candidates, 68 have repeated regional-variant support; none has support under the retained *≤*1% variant-frequency rule and none has RTT-variant support. Exact directed-candidate counts, including zero, are tabulated. Twenty-six source states receive positive-only ordering among existing ties. Auxiliary evidence ranks existing tied candidates; spacer membership and the spatial/RNA test cohorts are defined independently. Data derive from one 4T1 tissue section (biological n = 1), with two technical ONT libraries. Tissue spots, read observations, pooled spot–UMI keys, sequence states and directed candidate edges are distinct units. Downstream variation is reported as auxiliary cDNA evidence. The tables retain separate regional denominators and directed-candidate counts. i, All 289 repeated state–region–variant support entries, comprising 275 PBS and 14 RTT entries, plotted by regional frequency and number of spacer states carrying the variant. These entries represent 39 distinct regional variants (33 PBS and six RTT); multiple entries for the same variant can overlap. Frequency denominators are 8,690 PBS and 9,741 RTT callable pooled keys. Point area retains the original mapping, 8 + 2√(supporting spots), and both axes are logarithmic. The 1% line is diagnostic and does not change core membership. One tissue section was assayed with two technical spatial libraries, barcode12 and barcode13. Capture spots, exact states, extraction calls and pooled spot–UMI keys remain distinct units.

**Figure S7.**
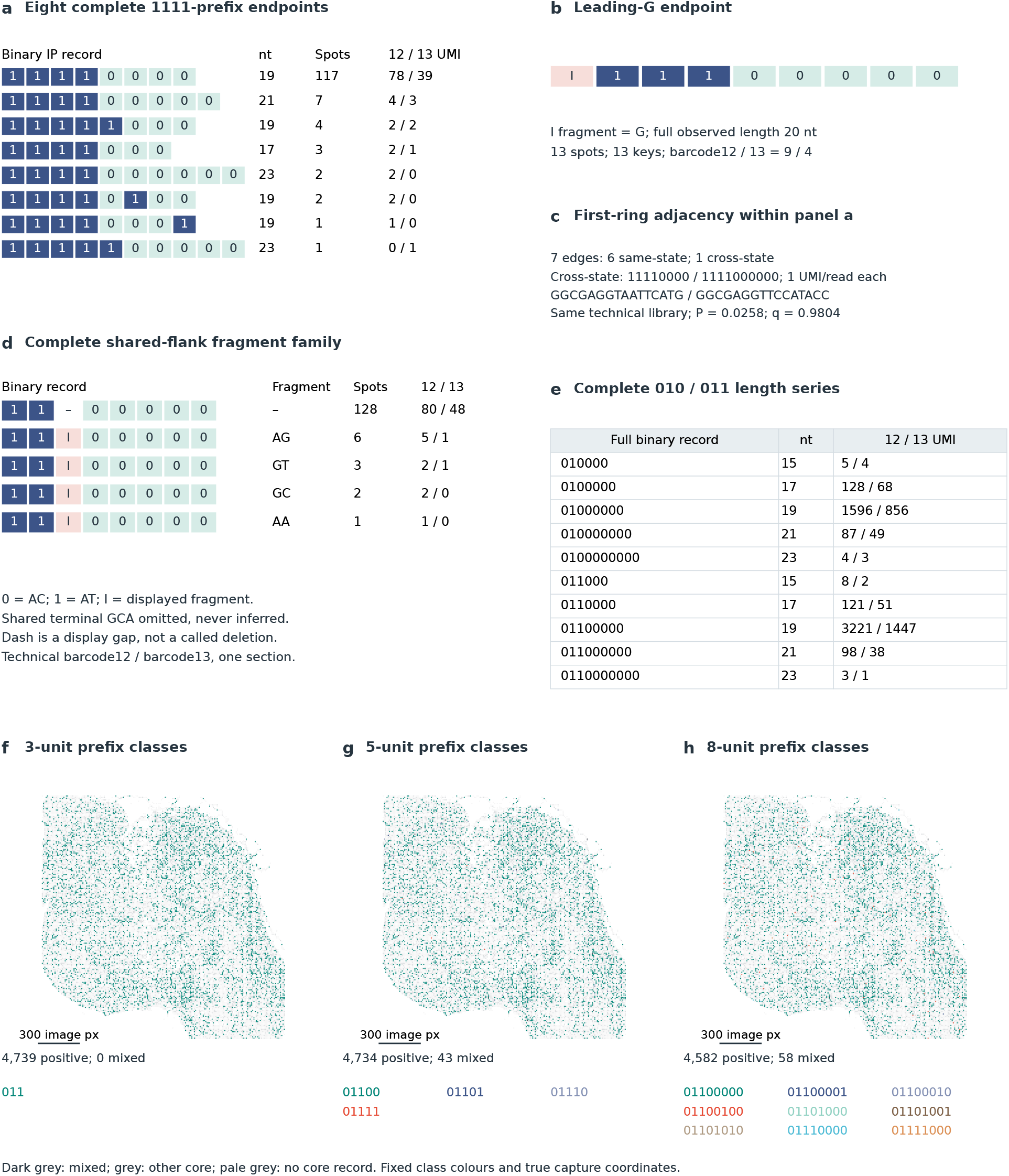
Complete spatial examples and nested prefix classes, related to Figures 5 and 6. a, Binary representation of all eight exact endpoints underlying Figure 6a,b, integrating complete records, full length and per-technical-library support. The eight states occupy 137 spots; the dominant endpoint occupies 117. Four endpoints are observed in both barcode12 and barcode13. Rows are ordered by pooled spot support. Exact sequence strings, stable IDs and each library’s counts are retained in the accompanying table. b, The leading-G endpoint underlying Figure 6c, with 13 pooled spots and 13 UMI keys, supported by nine and four keys in the two technical libraries. c, Seven first-ring edges within that history: six connect the same exact state and one connects different exact states. The cross-state endpoints each have one UMI and one read in the same technical library. The group-level test gives P=0.0258 and q=0.9804; edge counts are descriptive. d, The complete shared-flank family underlying Figure 5g: the canonical comparator and AG, GT, GC and AA fragments, at 128, 6, 3, 2 and 1 spots, respectively. Exact fragments accompany uniform indel display cells. e, Per-library support for all ten states in the 010/011 length series in Figure 5e. The complete binary record and full nucleotide length are shown together with the two technical-library counts. AC=0 and AT=1; the common terminal GCA is omitted. I marks the displayed noncanonical fragment, whose exact bases accompany the record. A comparator dash is a display gap, not a called deletion. f–h, The 011-class three-, five- and eight-unit prefix classes at actual spot centres, using hexagonal glyphs and common limits. There are 4,739, 4,734 and 4,582 focus-positive spots, including 0, 43 and 58 spots with multiple displayed classes. Mixed spots have a dedicated colour rather than a forced dominant label. Other core-bearing and core-unobserved tissue spots are retained as distinct grey backgrounds. Prefix classes were fixed using sequence and molecular support before spatial or RNA outcomes were examined. The paired model test is shown separately in Figure S10c. The complete 37-record matrix and three-, five- and eight-unit class-membership strips are shown in Figure S8d; exact sequences, masks and class assignments are supplied as Source Data. Maps retain every measured tissue position and use the display radius, colours, mixed-category definition and common limits. The ruler denotes 300 image pixels.

**Figure S8.**
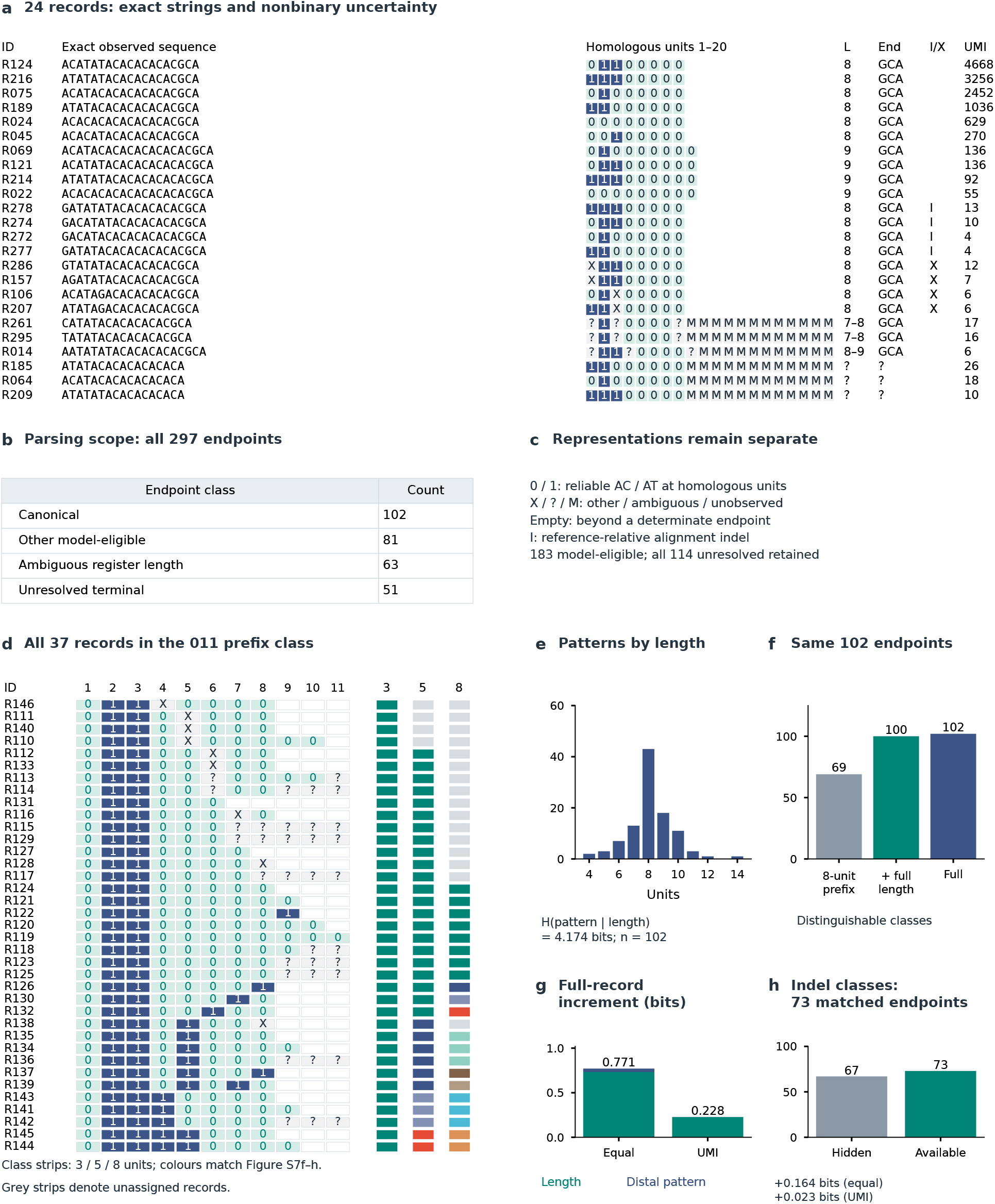
Complete record representation and information, related to Figures 3, 5 and 6. a, Twenty-four observed records selected before spatial analysis using fixed sequence categories and pooled molecular support: six canonical records of at most eight units, four extended canonical records, four supported structural variants, four records with other substitutions, three ambiguous-length records and three terminal-unresolved records. Display IDs map to stable source IDs and exact strings. Reliable AC and AT units are pale-teal 0 and navy 1. X denotes a nonbinary substitution,an ambiguous aligned position and M an unobserved position where the endpoint is not determinate. An empty outlined cell is outside a determinate record. L is the inferred register length or set of equally optimal lengths; End is recovered GCA or unresolved. I denotes a supported reference-relative insertion/deletion feature. UMI counts are distinct pooled spot-UMI keys. b, Parsing categories for all 297 exact endpoints: 102 canonical, 81 other model-eligible, 63 with ambiguous register length and 51 with unresolved terminal. The 183 model-eligible records form the shared model-comparison cohort. c, Representation key and analysis scope. The complete 297-record encoded ledger, exact strings, alignment alternatives and matrix are supplied as data files. The ledger also includes records with unresolved length or terminal status. d, All 37 endpoints in the 011 prefix class with binary characters, unresolved positions and class-membership strips for three-, five- and eight-unit views. AC=0, AT=1, X is another substitution,is unresolved and empty cells are beyond a determinate endpoint. I denotes a supported reference-relative indel feature. Exact sequences and masks remain in Source Data. The class-strip colours match the maps in Figure S7f–h; grey strips denote unassigned records. e, Number of distinct canonical binary patterns at each fixed register length in the same 102-endpoint spatial cohort. Conditional entropy given length is shown with equal endpoint weights; molecule-weighted distributions are provided in Source Data. f, Distinguishable classes under an eight-unit visible prefix, prefix plus explicitly restored full length, and complete record. Full length is included in the secondary contrast and omitted from the prefix-only projection. g, Partition of the entropy increment over the prefix into length and distal pattern contributions, separately using equal endpoint and pooled spot-UMI weights. The contrasts compare retained sequence content under the stated projections. The increment is conditional on the prefix: equal-endpoint full information is 6.672 bits, of which the prefix already contains 5.902 bits. Length and distal pattern add 0.732 and 0.039 bits, respectively. Full-precision values and the separate pooled-UMI decomposition are in Source Data; visible boundaries of shorter records are retained in the prefix. h, Exact structural identity hidden or available in the same 73 supported endpoints, with shared lengths and homology masks. The comparison isolates the incremental contribution of reliable exact structural identity. Information contrasts quantify endpoint discrimination within each specified cohort, using separate equal-endpoint and pooled-UMI weights.

**Figure S9.**
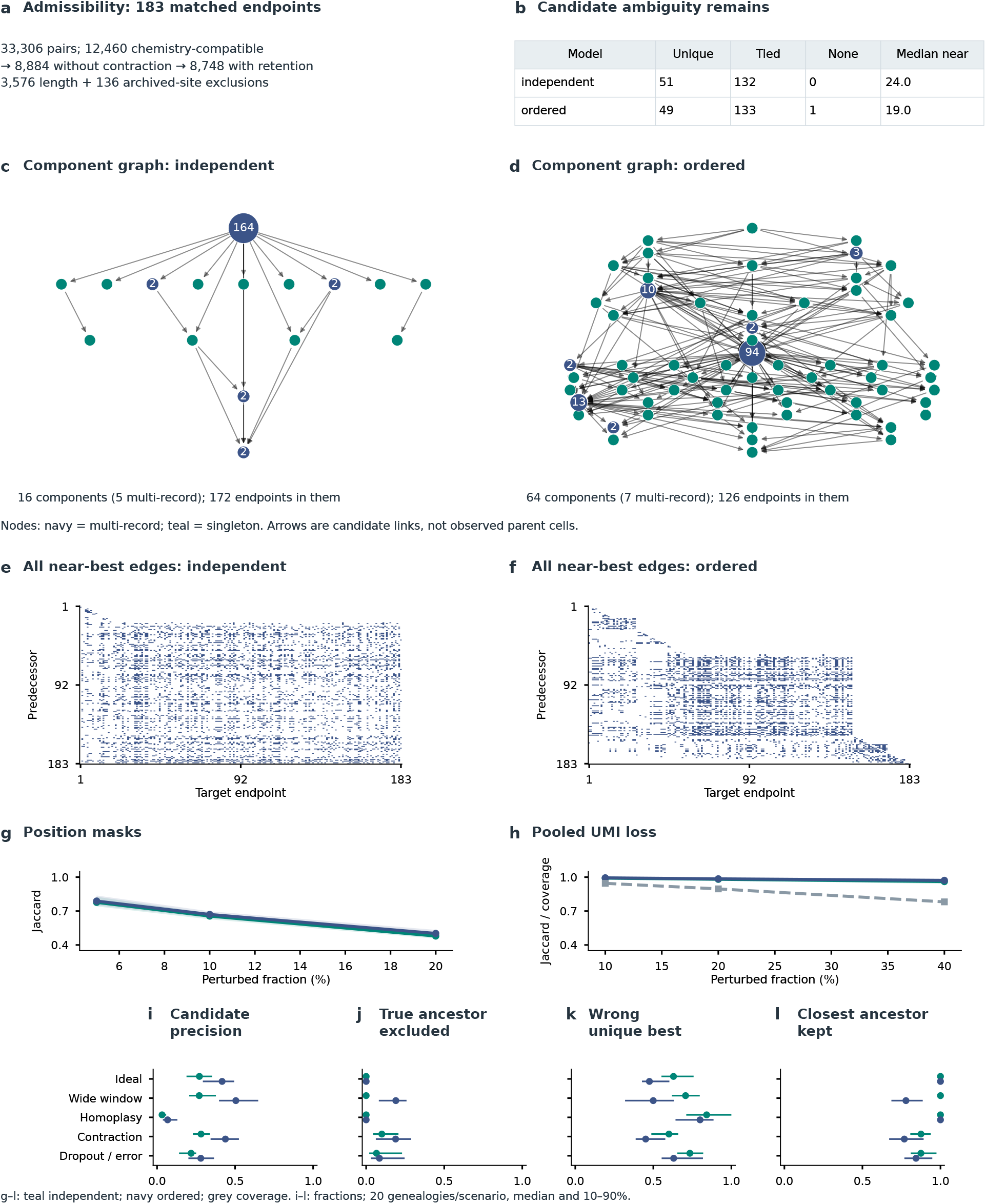
Candidate relations, stability and model mismatch, related to Figures 5 and 6. a, The same 183 records and 33,306 directed pair opportunities under shared chemistry, an added no-contraction constraint and an added archived-site retention rule. The exploratory retention window is eight units. Length exclusion removes 3,576 pairs, and archived-site retention removes 136 additional pairs. b, Endpoint counts with a unique best candidate, tied best candidates or no admissible candidate; the table gives the median near-best candidate count. Costs and the one-unit near-best margin are shared. Counts summarize candidate resolution under each model. c,d, Complete near-best relation graphs condensed at strongly connected components for the independent and ordered models. All 183 eligible endpoints are represented, including isolated components. Component size scales node area; labels give counts where more than one endpoint is present. The independent graph has 16 components, containing 172 endpoints in multi-record components; the ordered graph has 64 components, with 126 such endpoints. Arrows connect components of candidate sequence relationships. e,f, All near-best directed candidate edges, reordered by component and stable source ID for inspection. Matrix row/column mappings, all individual candidates and complete quotient edges are supplied. Graphs preserve tied relations, unplaced endpoints and strongly connected components. Records outside this comparison are retained in the complete 297-record data matrix. g, Candidate-set Jaccard after random masking of reliable positions, preserving their order. h, Conditional Jaccard among retained records after random pooled-UMI loss, with record coverage shown separately. Lines are medians and shading 10th-90th percentiles over 50 replicates per setting. The loss analysis holds state calls and feature definitions fixed. i–l, Separate known-genealogy simulations: near-candidate precision, fraction of true sampled-ancestor relations excluded, wrong-relation fraction among unique-best targets, and retention of the closest sampled ancestor. Points and bars are medians and 10th-90th percentiles across 20 independently seeded genealogies for each scenario. Five prespecified scenarios cover ideal terminal editing, a wider true window, saturation/homoplasy, contraction/re-editing and dropout/error. Both models receive identical observations. Unique-best status denotes score resolution. The root remains in overall target-coverage and unique-best denominators; ancestor-retention denominators contain only actual true relations. Simulation results evaluate the two models against the saved latent genealogies. In g–l, teal denotes the independent model and navy the ordered model; grey in h gives coverage. In c,d, navy denotes multi-record components and teal singleton components. The node coordinates follow the retained display layout, using the exact component membership and quotient-edge tables.

**Figure S10.**
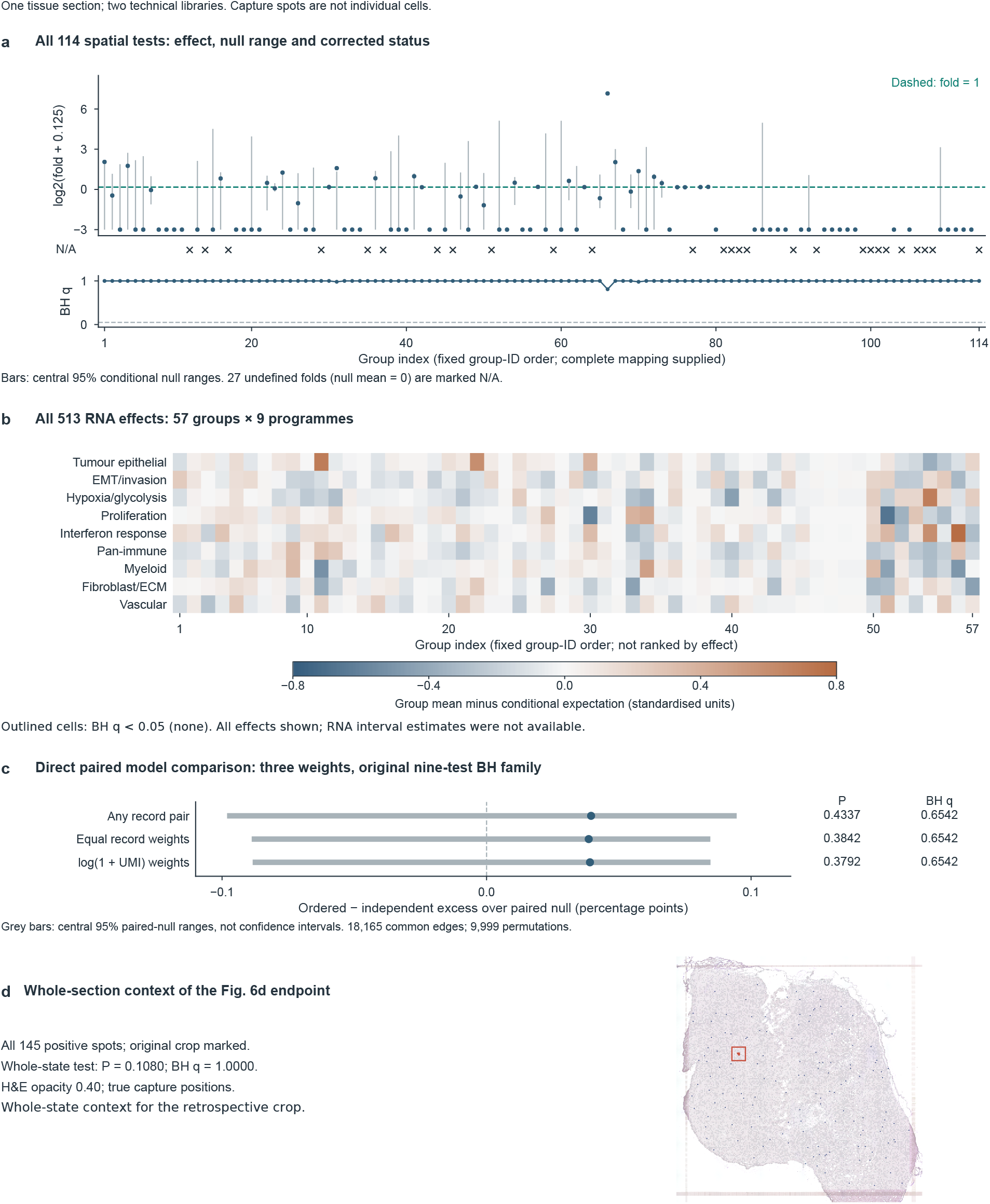
Association tests and whole-section context, related to Figure 6. One 4T1 tumour section was profiled with two technical ONT libraries (barcode12 and barcode13). Capture spots, not cells, are the spatial observation units. Groups can overlap in membership. a, All 114 spatial tests in fixed group-ID order. Points show log2(observed first-ring edges / conditional-null mean + 0.125); grey bars are the central 95% conditional permutation-null ranges divided by the same nonzero null mean and transformed identically. The dashed no-enrichment reference is log2(1.125), not zero. For 27 groups the null mean and observed edge count are both zero, so fold enrichment is undefined; these groups are marked N/A, not assigned zero effect. All 114 BH q values are shown below; the dashed threshold is q=0.05. The family uses 11,330 core-positive capture spots and 18,633 undirected first-ring edges, with 9,999 conditional permutations and no refined tests. Source Data for Figure S10 retains each group’s member count, raw edge count, null mean/range, exact P/q and membership. b, All 57 group-by-nine-programme comparisons (513 tests), with columns in fixed group-ID order and programme identities retained. Colour is the group mean minus its conditional expectation in standardised programme units; the symmetric -0.8 to +0.8 scale covers every effect without clipping. Outlines denote BH q<0.05; none is present. RNA P/q values are retained in Source Data for Figure S10 under their original 513-test family. RNA interval estimates were not available. Fifty-seven additional spatial groups failed the pre-existing RNA group-size eligibility rule; their status is documented separately and they are not added to the RNA BH family. c, Direct ordered-minus-independent comparisons under presence, equal-record and log(1+UMI) weighting on the same 18,165 eligible spatial edges. Points are the observed paired difference minus its paired-null mean, multiplied by 100; grey bars are the central 95% paired-null bounds centred on the same mean and multiplied by 100. For the primary presence measure, the uncentred observed difference is -0.369 percentage points and the excess over its paired-null mean is +0.040 percentage points. These are permutation-null ranges, not confidence intervals. The three displayed differences remain within the nine-test BH family, which also includes the six model-specific tests retained in Source Data for Figure S10. There were 9,999 paired whole-spot permutations. No spatial or RNA association, or paired model advantage, passed its corresponding correction family; non-significance does not establish absence or equivalence. d, Whole-section positions of all 145 spots carrying the complete state shown in the retrospective three-spot crop in Figure 6d. The crop location is retained. Whole-state P=0.1080 and BH q=1.0000; the map provides whole-state spatial context.

**Table S1.** Editor-analysis strata, related to Figure S4. The editor comparison uses the 19-nt starting-register reference (AC)8GCA (ACACACACACACACACGCA). The table lists the primary analysis strata. Counts are read occurrences. AC8 conversion uses the substitution-compatible 19-nt subset; same-length diversity uses the all-letter 19-nt subset. Complete sequence-class inventories and exact reference-sequence frequencies are provided in Source Data. Site-specific thresholds apply only to the focal site and include every allele there. Full-body thresholds apply at all positions, including unchanged bases. The five joint classes are mutually exclusive; conversion-event counts overlap because the “both” class contributes to both A-to-G and C-to-T. Strings in Other can also carry a target-compatible base alongside incompatible changes; their counts are tabulated separately from substitution-compatible numerators. Depth fractions were 0.05, 0.10, 0.25, 0.50, 0.75 and 1.00 of common depth. Each fraction used 200 without-replacement hypergeometric draws, seed 20261012. At common depth 53,089, NG-ABE8e had 107 strings and D1=3.940, with no sampling interval because the complete stratum was used. AncBE4max had mean richness 72.3 (technical 2.5–97.5% range 65–78) and D1=2.920 (2.886–2.958). These intervals do not quantify between-culture biological uncertainty. Minimum-support sensitivities remained separate from the primary singleton-retaining distributions.

| Analysed layer | NG-ABE8e occurrences | AncBE4max occurrences |
| --- | --- | --- |
| All admitted full-body Q20 observations | 109,964 | 905,949 |
| All-letter 19-nt bodies (AC8 length), full-body Q20 | 53,089 | 393,868 |
| AC8-substitution-compatible 19-nt bodies | 52,623 | 389,281 |

