## Supplementary figures and images for "Self-extending DNA recording with stochastic and programmed writing"

### tissue_image.png

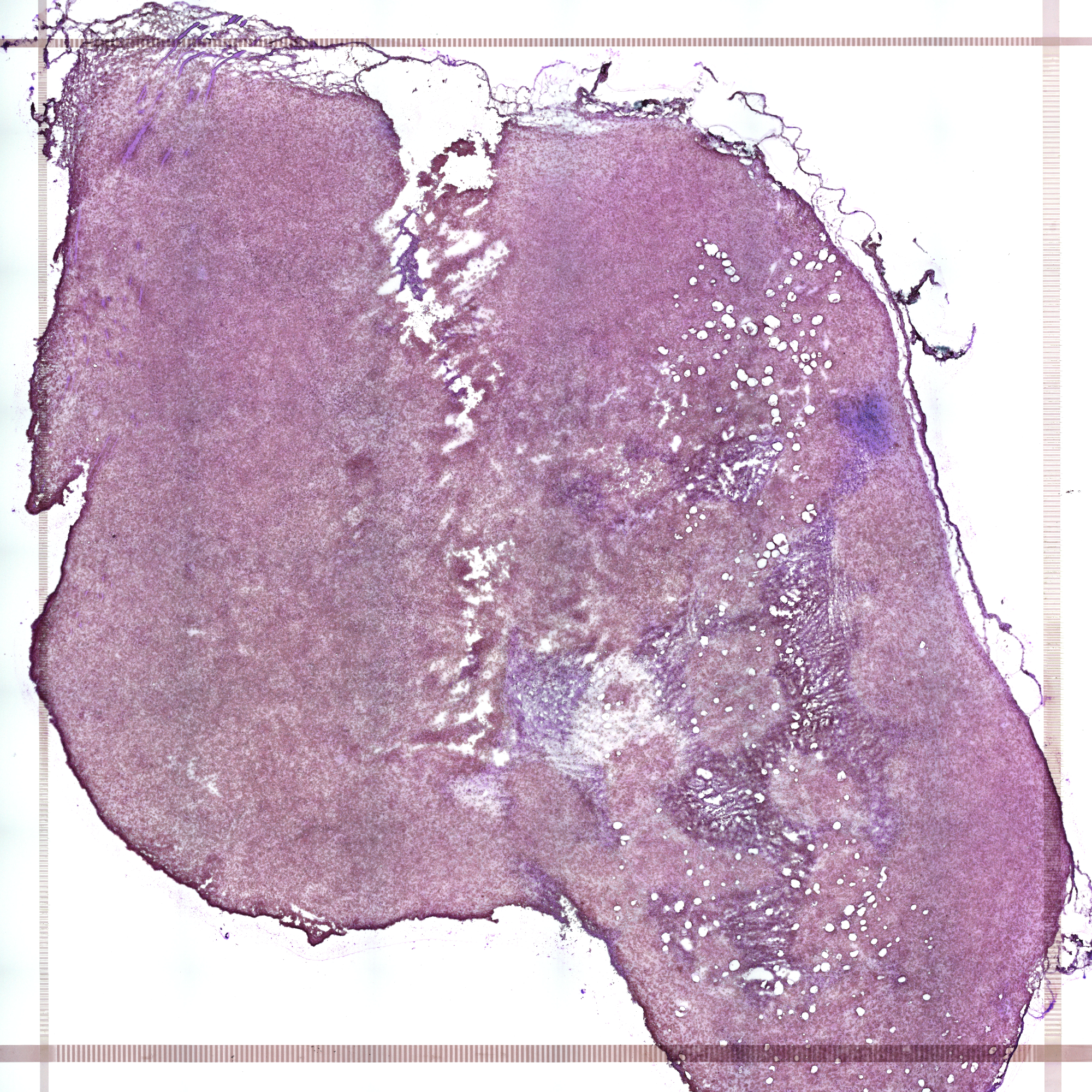
